# A critical role of FAK signaling in Rac1-driven melanoma cell resistance to MAPK pathway inhibition

**DOI:** 10.1101/2024.12.22.629990

**Authors:** Jesse D. Riordan, Teagan A. Nathanson, Afshin Varzavand, Adeline A. Hawkins, Rebekah M. Peplinski, Elizabeth C. Hannan, Faith A. Bibeau, Nathaniel J. Freesmeier, Madison C. Jilek, Silvia Coma, Jonathan A. Pachter, Adam J. Dupuy, Christopher S. Stipp

## Abstract

The Rac1 P29S hotspot mutation in cutaneous melanoma is associated with resistance to MAPK pathway inhibitors (MAPKi) and worse clinical outcomes. Moreover, activation of Rac1 guanine exchange factors (GEFs) also promotes MAPKi-resistance, particularly in undifferentiated melanoma cells. Here we delineate mechanisms of Rac1-driven MAPKi-resistance and identify strategies to inhibit the growth of this class of cutaneous melanomas. We find that Rac1-driven melanomas manifest pleiotropic resistance mechanisms including (i) reduced dependence on BRAF/MEK, (ii) activation of alternative MAPK pathways utilizing Jun kinase and p38 MAP kinase, and (iii) a partial reliance on YAP/TAZ signaling. Importantly, although Rac1-driven melanoma cells display reduced dependence on BRAF/MEK, they are not completely ERK-independent. Additionally, the presence of activated Rac1 appears to create a dependency on focal adhesion kinase (FAK) signaling in undifferentiated melanoma cells. Therefore, despite the pleiotropic mechanisms of Rac1-driven MAPKi resistance, we find that combined inhibition of RAF and MEK with the RAF/MEK clamp auvotometinib and FAK with the FAK inhibitor defactinib is a promising approach for suppressing the growth of Rac1-driven melanoma cells. Thus, the avutometinib plus defactinib combination, which is currently being investigated for brain metastatic cutaneous melanoma may also have utility against Rac1-driven MAPKi-resistance in heavily pre-treated, advanced disease.

## Introduction

Mutations in the monomeric GTPase gene RAC1 are the third most common hotspot mutation in melanoma after BRAF and NRAS mutations (1). Such mutations render the Rac1 protein constitutively active, resulting in unscheduled signaling towards multiple effectors, including PAK kinases, Arp2/3, AKT, and the SRF/MRTF transcriptional complex (2–9). In cutaneous melanoma, the most prevalent activating RAC1 mutation is a point mutation resulting in the protein change of P29S, which results from a C>T nucleotide substitution associated with UV damage (1). However, as visualized by cBioportal (10, 11), cancer genome sequencing reveals additional RAC1 hotspot mutations scattered across many different cancer types (**Table S1**). In melanoma, activating Rac1 mutations appear not to function as primary driver mutations; however, they may cooperate with driver mutations to promote melanoma cell proliferation, tumor progression, and metastasis, resistance to targeted therapy, and worse patient outcomes (6, 9, 12, 13).

Monomeric GTPases such as Rac1 are activated by guanine exchange factors (GEFs), and we previously identified upregulated DBL family member GEFs, including Vav1, as drivers of resistance to BRAF inhibitors (BRAFi) in a forward genetic screen using BRAF-mutant melanoma cells (14). However, the requirement of Rac1 for GEF-dependent BRAFi-resistance was not firmly established, nor was the mechanism by which Rac1 might drive BRAFi resistance downstream of DBL family GEFs. Vav1 is regulated by Src family tyrosine kinases, and co-targeting of BRAF and Src appeared to be a promising strategy for overcoming GEF/Rac1-driven BRAFi resistance (14). However, selective Src inhibitors are no longer being developed as clinical anti-cancer therapies.

Here we sought to elucidate the mechanisms by which DBL family GEFs and Rac1 can drive BRAFi resistance and identify additional strategies for targeting Rac1-driven drug-resistant melanomas. We confirmed a critical role for Rac1 in BRAFi resistance conferred by Vav1 and identified complex, pleiotropic mechanisms by which Rac1 and Rac1 GEFs can drive BRAFi resistance in melanoma. The Rac1-driven resistance mechanisms involve reduced dependence on canonical ERK pathway signaling and contributions from multiple additional signaling inputs. Despite the pleiotropy of the Rac1-driven resistance pathways, we identified focal adhesion kinase (FAK) inhibition as a promising strategy for the treatment of Rac1-driven drug-resistance in melanoma.

## Results

### Rac1 is required for Vav1-driven BRAFi resistance in V600-mutant melanoma cells

We recently found that upregulation of DBL family guanine exchange factors (GEFs), such as Vav1, can promote BRAFi/MEKi resistance in BRAF V600-mutant melanoma cells (14). It has previously been shown that Vav1 might signal through Rac1, RhoA, or Cdc42 (15). Our initial study suggested that Vav1 might promote BRAFi resistance specifically by enhancing Rac1 activity (14); however, whether Rac1 was required for Vav1-driven BRAFi resistance had not been established. Therefore, we first created subclones of A375 melanoma cells with stable shRNA-mediated Rac1 knockdown (Rac1 KD) or cells harboring a non-targeting shRNA (NT). We then transduced the Rac1 KD and NT subclones with either a VAV1 expression vector (VAV1) or an empty vector control construct (EV). Overexpression of Vav1 in A375 NT subclones F2 and B7 clearly promoted BRAFi-resistance; however, Vav1 over-expression was unable to promote BRAFi resistance in Rac1 KD subclones G3 and F10, in which Rac1 was stably knocked down by RNAi (**Fig. 1A, B**). As expected, the empty vector control construct did not promote BRAFi resistance in either Rac1 KD or NT subclones (**Fig. 1A,B**). Vav1 over-expression had minimal effects on cell proliferation in the absence of BRAFi, whether Rac1 was knocked down or not (**Fig. 1C, D**). Immunoblotting confirmed strong knockdown of Rac1 in the Rac1 shRNA cells and increased Vav1 expression in the Vav1-overexpressing subclones (**Fig. 1E**). These data indicated that Rac1 is required for Vav1-mediated BRAFi resistance.

**Figure 1.**
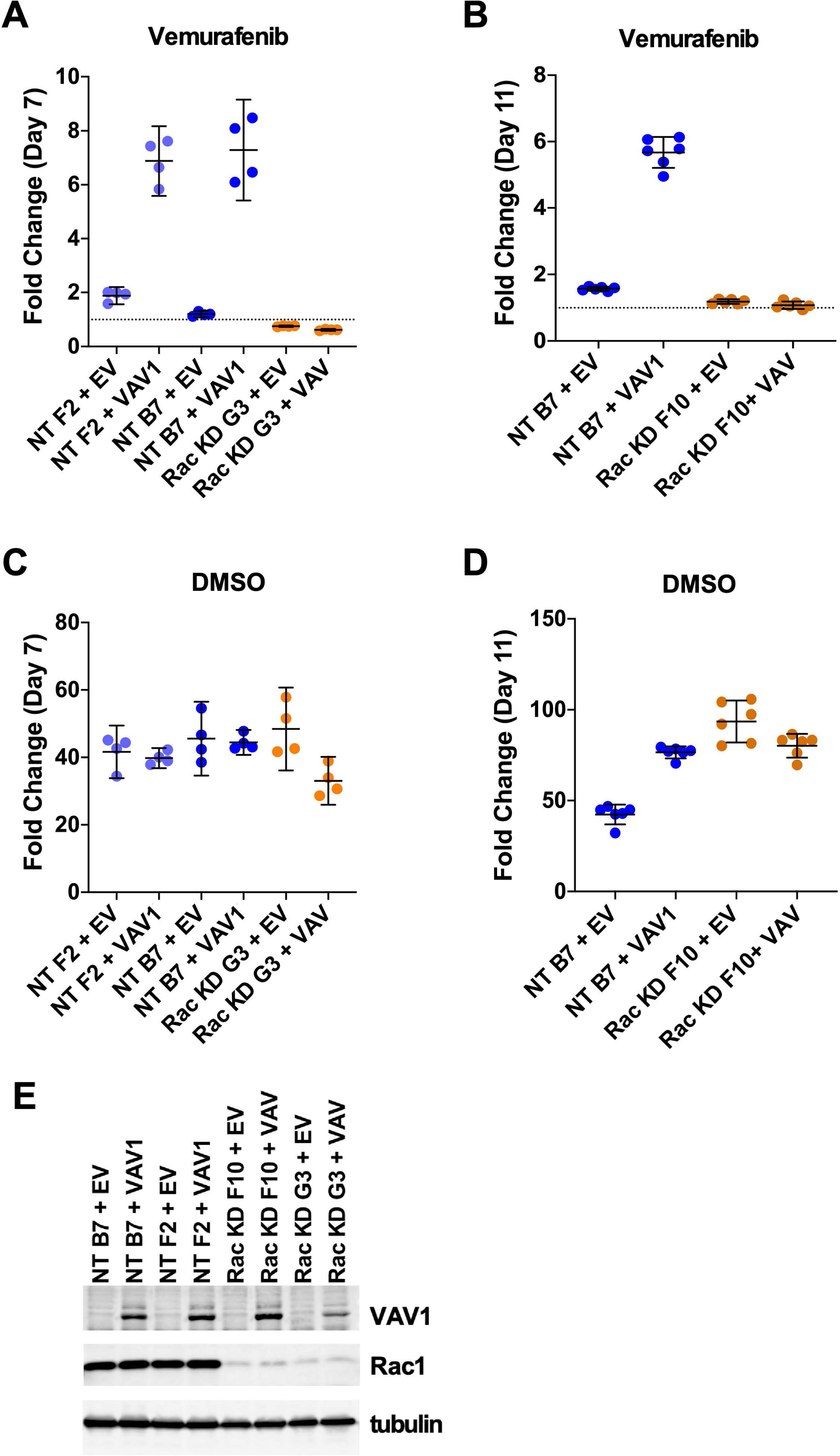
Rac1 is required for Vav1-driven BRAFi resistance in BRAF-mutant melanoma. **(A)** BRAF V600E mutant A375 subclones NT F2 and NT B7 harbor a non-targeting control shRNA for Rac1, compared to A375 subclone G3, which harbors an effective shRNA targeting Rac1. Cells were grown in 3 µM vemurafenib (**VEM**). Over-expression of Vav1 in the NT F2 and NT B7 subclones (**NT F2 + VAV1**, and **NT B7 + VAV1**), but not the introduction of an empty vector control **(NT F2 + EV**, and **NT B7 + EV**) promoted resistance to the BRAFi vemurafenib (blue symbols). In contrast, in A375 subclone G3, which harbors an effective Rac1 shRNA, Vav1 over-expression was unable to promote vemurafenib resistance compared to empty vector control cells (**Rac KD G3 + EV** vs. **Rac KD G3 + VAV1**; orange symbols). **(B)** In a separate experiment, similar to panel A, Vav1 over-expression in A375 subclone B7 (with a non-targeting control shRNA for Rac1) promoted vemurafenib resistance (**NT B7 + EV** vs **NT B7 VAV1**; blue symbols), while over-expression of Vav1 in a different subclone F10, which harbors an effective Rac1 shRNA, was unable to promote vemurafenib resistance, compared to Vav1 empty vector control cells (**Rac KD F10 + EV** vs **Rac KD F10+ VAV1**; orange symbols). **(C & D)** Regardless of Rac1 knockdown or Vav1 over-expression status, all the sublines showed similar high levels of growth in experiments using DMSO vehicle control instead of the BRAFi, vemurafenib. **(E)** Immunoblotting confirmed that Rac1 was knocked down in the F10 and G3 A375, compared to the F2 and B7 subclones, and that Vav1 protein was over-expressed in the Vav1 (VAV1) cell line variants compared to their respective empty vector (EV) controls.

### PAK kinase signaling can contribute to Rac1-driven BRAFi resistance in BRAF V600-mutant melanoma cells but is not required for drug-resistant cell survival in the absence of BRAF inhibitors

It has been previously shown that some BRAFi/MEKi-resistant BRAF V600-mutant melanoma cells strongly upregulate PAK signaling and become dependent on PAK signaling for survival to varying degrees, such that certain pan-PAK inhibitors may be cytotoxic to BRAFi/MEKi-resistant cells in the absence of BRAF inhibitors (5). In contrast to these earlier results, we found that the PAK1-selective inhibitor G-5555 (16) partially suppressed the BRAFi-resistant phenotype of Rac1-driven drug-resistant melanoma cells in the presence of BRAFi treatment (vemurafenib; VEM) (**Fig. 2A, B**), but had no effect on cell survival in the absence of BRAF treatment (**Fig. S1**). Immunoblotting confirmed that G-5555 abolished PAK signaling towards MEK, almost completely eliminating phospho-MEK S298, a PAK-controlled phosphorylation site on MEK, in either the presence or absence of BRAFi treatment (**Fig. 2C**).

**Figure 2.**
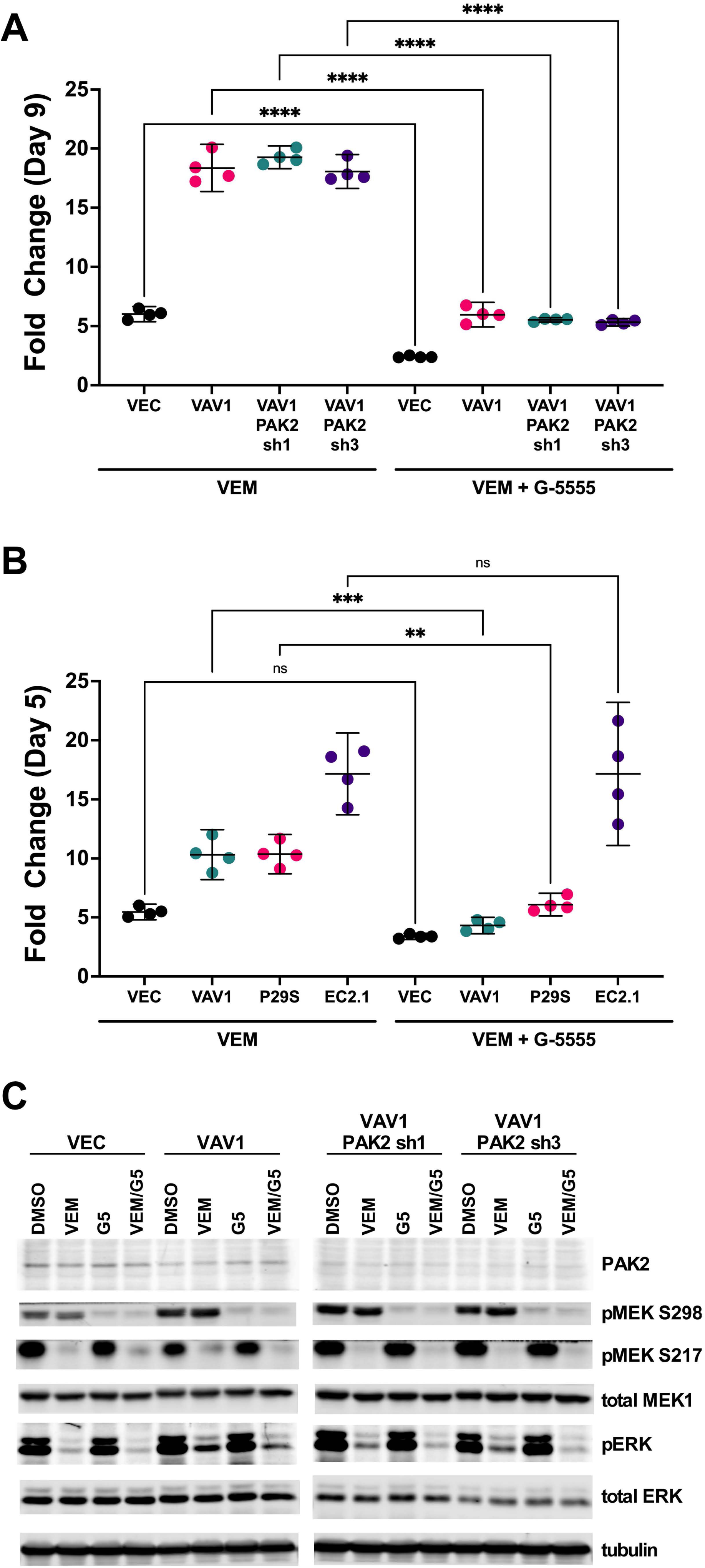
PAK kinase contributes to Rac-driven BRAFi resistance, but not to BRAFi resistance driven by a BRAF truncation. **(A)** The graph shows the fold change in A375 melanoma cell growth measured on day 9 by resazurin assay for vector control (**VEC**), Vav1 overexpressing (**VAV**), and Vav1 overexpressing cells with PAK 2 knock down by two different shRNAs (**VAV1 PAK2 sh1** and **VAV1 PAK2 sh3**). Cells were grown in 3 µM vemurafenib (**VEM**) or 3 µM vemurafenib plus 1 µM G-5555, a PAK1-selective inhibitor (**VEM + G-5555**). Vav1 overexpression promoted resistance to vemurafenib, which was significantly suppressed by G-5555. Similar results were observed in Vav1 overexpressing cells with PAK2 knockdown. **** P<0.0001, ANOVA with Sidak’s multiple comparison test. **(B)** The graph shows the fold change in A375 melanoma cell growth measured on day 5 by resazurin assay for vector control cells (**VEC**), Vav1-overexpressing cells (**VAV1**), Rac1 P29S expressing cells (**P29S**), and cells harboring a spontaneously acquired BRAF truncation (**EC2.1**). Cells were grown in 3 µM vemurafenib (**VEM**) or 3 µM vemurafenib plus 1 µM G-5555, a PAK1-selective inhibitor (**VEM + G-5555**). Inhibition of PAK1 by G-5555 significantly blocked vemurafenib resistance in Vav1 and Rac1 P29S-expressing cells, but not in EC2.1 cells with a BRAF truncation. *** P<0.001, **P<0.01, ANOVA with Sidak’s multiple comparison test **(C)** A375 vector control cells (**VEC**), Vav1 overexpressing cells (**VAV1**), and Vav1 overexpressing cells with PAK 2 knock down by two different shRNAs (**VAV1 PAK2 sh1** and **VAV1 PAK2 sh3**) were treated overnight with DMSO vehicle (**DMSO**), 3 µM vemurafenib (**VEM**), 1 µM G-5555 (**G5**), or 3 µM vemurafenib plus 1 µM G-5555 (**VEM/G5**). Cell lysates were then analyzed by SDS-PAGE and immunoblotting for PAK2, phospho-MEK S298, phospho-MEK S217, total MEK1, phospho-ERK, total ERK, and a tubulin loading control. Phospho-MEK S298 appeared increased in VAV1 over-expressing cells, compared to vector control cells, consistent with increased Rac1/PAK signaling. Treatment with G-5555 almost completely blocked phospho-MEK S298 in all four cell types.

In A375 melanoma cells, PAK2 appears to be a minor component of overall group I PAK expression (**Fig. 2C**), and silencing PAK2 by RNAi did not enhance the effect of the PAK1-specific G-5555 inhibitor (**Fig. 2A, B**). The remaining group 1 PAK, PAK3, was not detectable in A375 cells by immunoblotting (data not shown). Thus, the relatively non-cytotoxic PAK1-specific inhibitor, G-5555, almost completely canceled PAK signaling towards MEK1/2, but it had no effect on its own in suppressing the proliferation of BRAF V600-mutant melanoma cells under standard growth conditions. However, G-5555 partially re-sensitizied Rac1-driven BRAFi-resistant tumor cells to BRAF inhibition, both for Vav1 over-expressing cells and for cells expressing the clinically relevant Rac1 P29S constitutively active mutant (**Fig. 2A, B**). In contrast, a BRAFi-resistant A375 subline expressing a BRAF truncation mutant (A375 EC2.1) was immune to the effects of G-5555 (**Fig. 2B**), consistent with the Rac1-independent resistance mechanism of truncated BRAF, which cannot be inhibited by FDA-approved BRAF inhibitors, such as vemurafenib (17, 18). Expressing Rac1 P29S in 451Lu cells, another BRAF V600-mutant human melanoma cell line, also promoted BRAFi-resistance that was partially dependent on PAK signaling, based on PAK1 inhibition by G-5555 (**Fig. S2A**). Again, G-5555 on its own had no cytotoxic effect (**Fig. S2B**), revealing a role for PAK1 specifically in the context of BRAFi resistance than can be separated from the effects of more generally cytotoxic of pan-PAK inhibitors (5). Immunoblotting confirmed that phospho-MEK S298 was increased in the Rac1 P29S 451Lu cells and that G-5555 almost completely blocked MEK S298 phosphorylation (**Figs. S2C**).

### The Vav1-driven BRAFi resistance mechanism is highly resilient to MEK1/2 inhibition

Because PAK contributed to Rac1-driven BRAFi resistance, and PAK phosphorylation of MEK at the S298 site has been proposed to promote activation of the MAPK pathway (19, 20), we directly tested the contribution of MEK1/2 to the Vav1-dependent drug resistance phenotype. First, we hypothesized that if PAK phosphorylation of MEK1/2 contributed to BRAFi resistance, then a phosphomimetic MEK mutant (S298D) could be sufficient to promote drug resistance. This mutant has previously been suggested to functionally mimic constitutive phosphorylation of MEK by PAK (21). However, expression of MEK S298D did not promote resistance to vemurafenib any better than did expression of wild type MEK (**Fig. S3**). In contrast, expression of MEK S217D/S221D, a phosphomimetic MEK mutant at the BRAF-controlled S217/S221 sites, promoted strong primary resistance to vemurafenib (**Fig. S3**). Thus, a MEK mutant intended to mimic constitutive phosphorylation by PAK was not sufficient to promote BRAFi resistance, although a MEK mutant intended to mimic constitutive phosphorylation by BRAF did promote BRAFi resistance.

To test the extent to which MEK1/2 were necessary for Vav1-driven BRAFi resistance, we combined strong RNAi depletion of MEK1 (individually or in combination with MEK2) with pharmacological inhibition of BRAF (vemurafenib) or BRAF plus MEK (vemurafenib/cobimetinib). Vemurafenib or vemurafenib/cobimetinib combined treatment strongly suppressed the proliferation of vector control cells (**Fig. 3A**), and as previously observed, Vav1 expression promoted a level of resistance to BRAFi/MEKi (**Fig. 3B**). Surprisingly, strong knockdown of MEK1, the major MEK isoform in A375 cells, had no negative effect on Vav1-dependent BRAFi/MEKi resistance. If anything, Vav1-expressing cells with MEK1 knockdown displayed even stronger BRAFi/MEKi resistance than Vav1 cells with wild type MEK1 expression (**Fig. 3C**, compare to **Fig. 3B**). Double MEK1/2 knockdown also failed to blunt Vav1-driven BRAFi/MEKi resistance (**Fig. 3D**), and as with MEK1 single knockdown, Vav-1-expressing cells with MEK1/2 double knockdown were, if anything, even more resistant to BRAFi/MEKi treatment than Vav1-expressing cells with wild type MEK1/2 expression (compare **Fig. 3D** to **Fig. 3B**). Immunoblotting confirmed strong knockdown of both MEK1 and MEK2, as well as enforced Vav1 expression in this cell panel (**Fig. 3E**). Immunoblotting of lysates recovered at the end of the cell proliferation assays confirmed that Vav1 over-expression and MEK1 and MEK2 knockdown remained stable during the experiment (**Fig. 3F**). Immunoblotting also revealed that activated ERK1/2 was strongly suppressed, but not completely extinguished, in BRAFi/MEKi-treated, Vav1-expressing cells in which MEK1/2 were knocked down by RNAi (**Fig. 3F**). Thus, Vav1-driven BRAFi/MEKi resistance is highly resilient to genetic depletion of MEK1/2.

**Figure 3.**
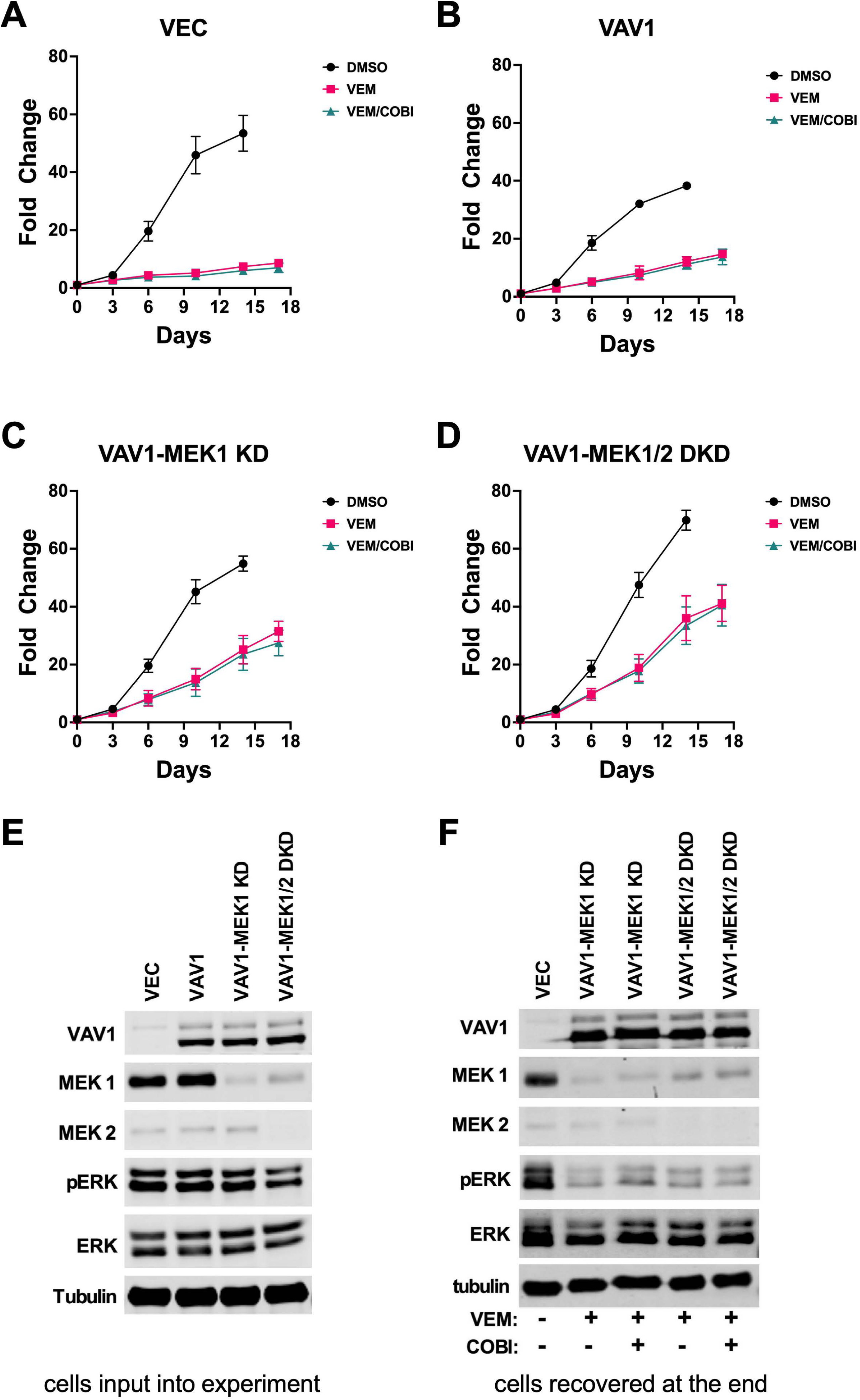
BRAFi-resistance resulting from Vav1 over-expression is highly resilient to MEK1/2 genetic depletion and pharmacological inhibition. (A-D) Vector control A375 cells (**VEC**), Vav1 overexpressing A375 cells (**VAV1**), Vav1 overexpressing A375 cells with MEK1 RNAi knockdown (**VAV1-MEK1 KD**), and Vav1 overexpressing A375 cells with MEK1 and MEK2 double knockdown (**VAV1-MEK1/2 DKD**) were grown in the presence of DMSO vehicle control, 3 µM vemurafenib (VEM), or 3 µM vemurafenib with 3 nM cobimetinib (VEM/COBI). Fold change growth in cell growth was measured over 15-18 days via resazurin assays. Knockdown or MEK1 or MEK1 together with MEK2 did not increase sensitivity of the cells to vemurafenib or vemurafenib/cobimetinib treatment. If anything, MEK1/2 knockdown cells grew better in the presence of BRAFi/MEKi treatment than Vav1 over-expressing cells without MEK1/2 knockdown. **(E)** Immunoblotting analysis of cells input into the experiment showed strong Vav1 overexpression in the Vav1 overexpressing cells and strong knockdown of MEK1 and MEK2 in the MEK1 and MEK1/2 double knockdown cells. **(F)** Immunoblotting of lysates from cells recovered at the end of the experiment showed that Vav1 over-expression and MEK1/2 knockdown were all maintained in the respective cell lines cultured under vemurafenib or vemurafenib/cobimetinib-treated conditions.

### A subset of spontaneously BRAFi/MEKi-resistant melanoma cells also depend on Rac1-PAK signaling and display resilience to MEK inhibition

We previously described the creation of a panel of spontaneously BRAFi/MEKi-resistant A375 melanoma cells that displayed multiple different resistance mechanisms arising within the same parental cell population (14). These resistance mechanisms include (i) rearrangements in the BRAF gene resulting in truncation of the N-terminal regulatory domain or tandem duplication of the BRAF kinase domain, (ii) loss of Ras-GAP NF1, and (iii) a slower arising form of drug resistance associated with a more spread cell morphology and increased phospho-MEK S298, a characteristic of cells with increased Rac1 signaling (Ref (14) and **Fig. S4A**). Three independently derived subpopulations with slower BRAFi-resistance kinetics were established: vemurafenib-resistant, persistent populations 1-3 (VRPP1, VRPP2, and VRPP3) (14, 22). RNAseq experiments revealed that VRPP1-3 have a distinct gene expression profile from all the other spontaneous resistance mechanisms, a profile that is defined in part by a YAP/TAZ-TEAD activation signature (23) (**Fig. S4B**). VRPP1 and VRPP2 cells are Rac1 wild type; however, VRPP3 cells acquired a Rac1 N92I mutation that went to fixation in the population, with an allele frequency of 1 mutant: 2 wild type copies of the Rac1 gene (22). Both the N92I and P29S mutations render Rac1 GEF-independent and therefore constitutively active (24, 25). Transient Rac1 siRNA experiments had suggested that the VRPP cell type depends on Rac1 for proliferation and drug resistance (22), an idea that we confirmed with strong, retrovirally delivered, shRNA-mediated Rac1 knockdown in VRPP3 cells (**Fig. S4C**). Moreover, although they were initially selected to be resistant to the single agent BRAFi, vemurafenib, the VRPP lines were also resistant to combination BRAFi/MEKi treatment. Consistent with our previous observations on the potential utility of combined BRAF/Src inhibition for overcoming Rac-driven BRAFi-resistance (14, 22), a Src family kinase inhibitor, saracatinib, resensitized the VRPP2 and VRPP3 cells to BRAF inhibition (**Fig. S4D**). Further analysis of the VRPP cell gene expression profile revealed increased expression of undifferentiated melanoma cell markers together with reduced expression of neural crest cell markers (26). This change in gene expression is consistent with a VRPP cell transition along the differentiation trajectory described by Tsoi et al. (26), from a neural crest-like state to an undifferentiated state as the cells acquired resistance to BRAF inhibition (**Fig. S5**)

Like A375 melanoma cells expressing Vav1 or Rac1 P29S, the BRAFi-resistance phenotype of VRPP1-3 cells was blunted by the PAK1 inhibitor G-5555 (**Fig. 4**), confirming that PAK1 can also contribute to spontaneous Rac1-driven BRAFi resistance. Like VAV1 overexpressing cells, Rac1 mutant VRPP3 cells displayed resilience to combined MEK1/2 RNAi depletion coupled with combined BRAFi/MEKi pharmacological inhibition (**Fig. 5**). Thus, a subset of spontaneously BRAFi-resistant V600-mutant melanoma cells also utilize a Rac1-dependent pathway to become drug resistant. In A375 melanoma cells, this spontaneous Rac1-driven resistance pathway is associated with an undifferentiated gene expression profile, activation of a YAP/TAZ-TEAD signature, and resilience to MEK1/2 inhibition and genetic depletion. Collectively, these results support the significance of upregulation of Rac1 signaling as an important mechanism of BRAFi/MEKi resistance.

**Figure 4.**
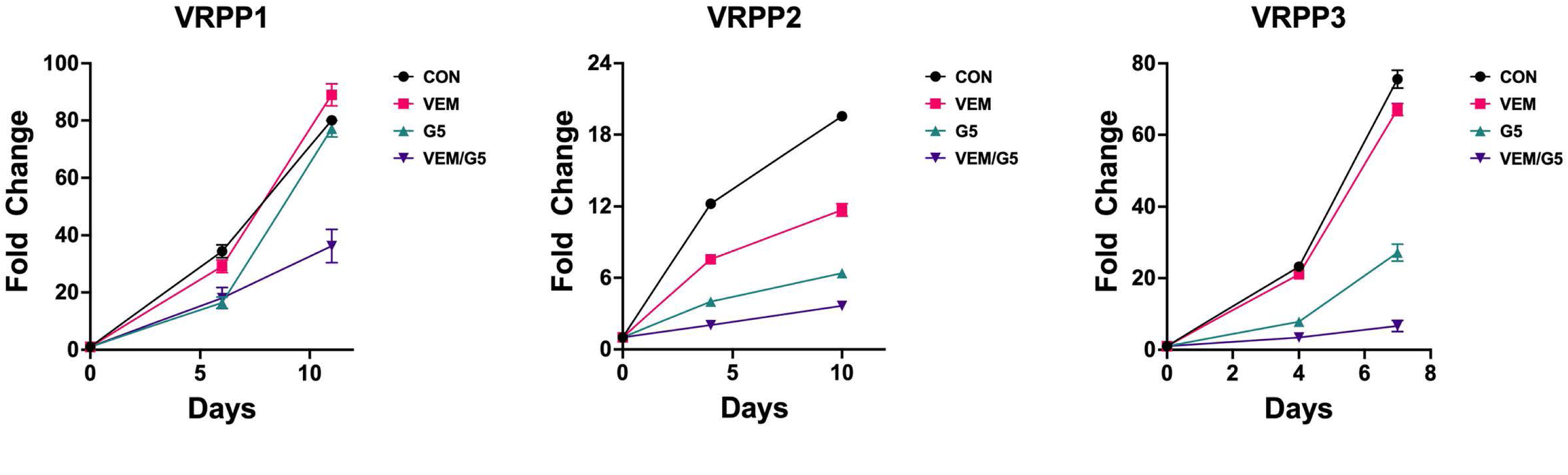
PAK signaling contributes to the spontaneous BRAFi resistance of the A375 VRPP cell lines. A375 VRPP1, VRPP2, and VRPP3 cell lines were grown in the presence of DMSO vehicle (**CON**), 3 µM vemurafenib **(VEM**), 1 µM G-5555 (**G5**), or 3 µM vemurafenib plus 1 µM G-5555 (**VEM/G5**), and fold change in cell growth was measured over time via resazurin assay. Co-treatment with G-5555 significantly reduced the resistance of the VRPP cell lines to vemurafenib.

**Figure 5.**
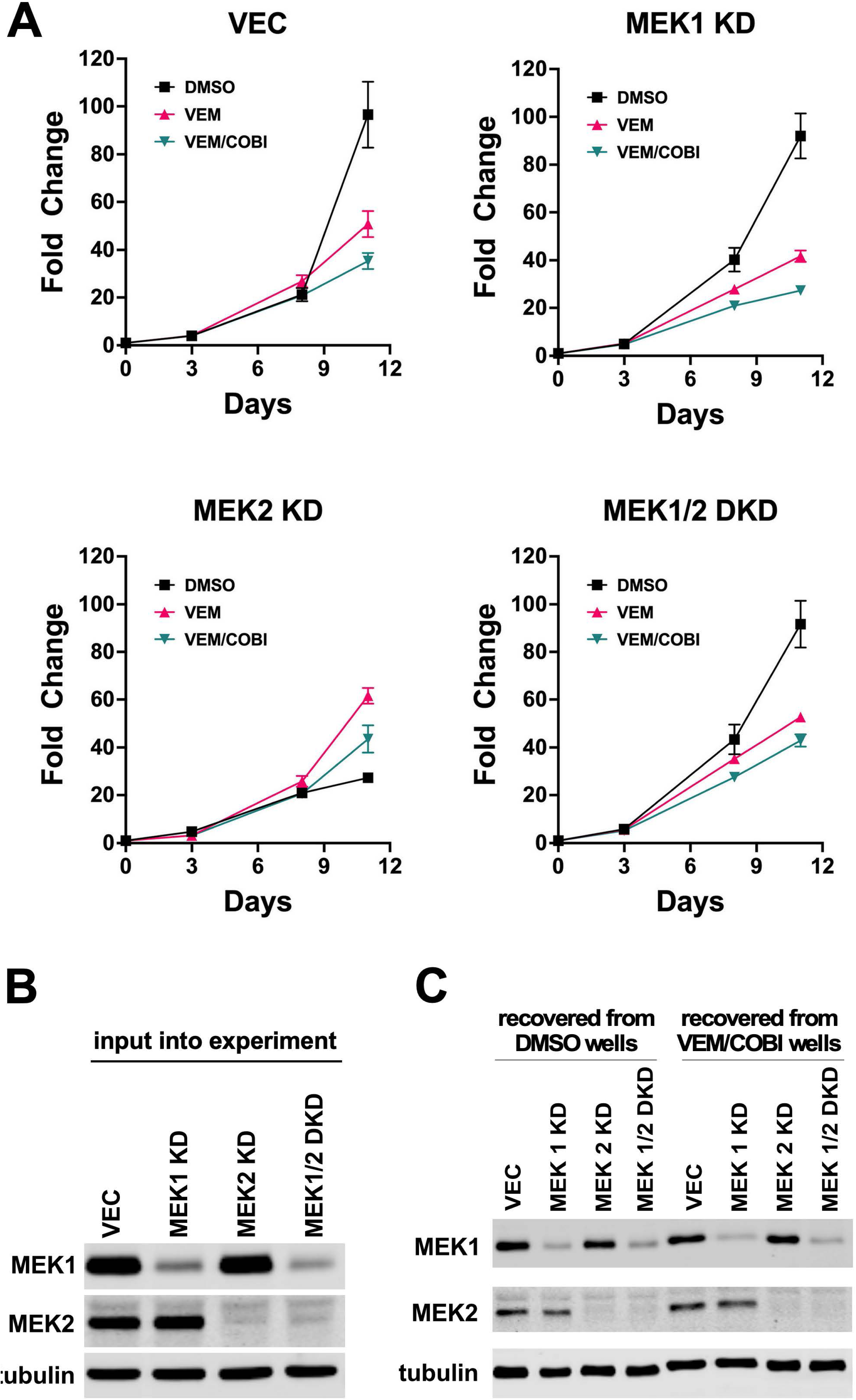
Spontaneously BRAFi-resistant A375 VRPP3 cells with a Rac1 N92I mutation are also highly resilient to MEK1/2 genetic depletion and pharmacological inhibition. **(A)** VRPP3 cells with non-targeting shRNAs (**VEC**) and cells with MEK1 knockdown (**MEK1 KD**), MEK2 knockdown (**MEK2 KD**), or MEK1/2 double knockdown (**MEK1/2 DKD**) were grown in the presence of DMSO vehicle, 3 µM vemurafenib (**VEM**) or 3 µM vemurafenib plus 3 nM cobimetinib (**VEM/COBI**). Fold change cell growth was measured over time via resazurin assay. All of the MEK knockdown cell lines grew comparably to vector control in the presence of vemurafenib or vemurafenib/cobimetinib treatment. SDS-PAGE and immunoblotting of lysates from the cell lines input into the experiment (B), and from cells recovered at the end of the experiment **(C)** confirmed strong, stable knockdown of MEK1 and MEK2 in the respective cell lines.

### Rac1-driven resistance mechanisms are partially but not completely resilient to ERK1/2 inhibition

Given the resilience of multiple Rac1-driven cells to BRAFi/MEKi treatment, we wondered whether these cells might be ERK-independent, as previously suggested for certain classes of Rac1/PAK-driven, BRAFi/MEKi-resistant melanoma cells (5). To test the requirement for ERK1/2 activity in Rac1-driven cells, we treated cells with different doses of the highly ERK1/2-selective inhibitor, ulixertinib. Compared to parental A375 cells, cells expressing Vav1, Rac1 P29S, or the spontaneously-resistance Rac1 N92I-expressing VRPP3 cells all displayed enhanced resistance to ulixertinib treatment (**Fig. 6**). In contrast, A375 EC2.1 cells, with a BRAF truncation resistance mechanism, were even more sensitive than parental A375 cells to ERK inhibition with ulixertinib (**Fig. 6**). In addition, 451Lu melanoma cells expressing Rac1 P29S were also somewhat more resistant to ERK inhibition compared to vector control cells (**Fig. S6**). However, all cell types tested were inhibited by higher doses of ulixertinib (**Figs. 6 & S6**). Thus, the Rac1-driven drug-resistant cells we have examined in detail thus far have displayed a striking resilience to BRAFi/MEKi treatment and a partial resilience to ERK1/2 inhibition, but they do not appear to be completely ERK-independent.

**Figure 6.**
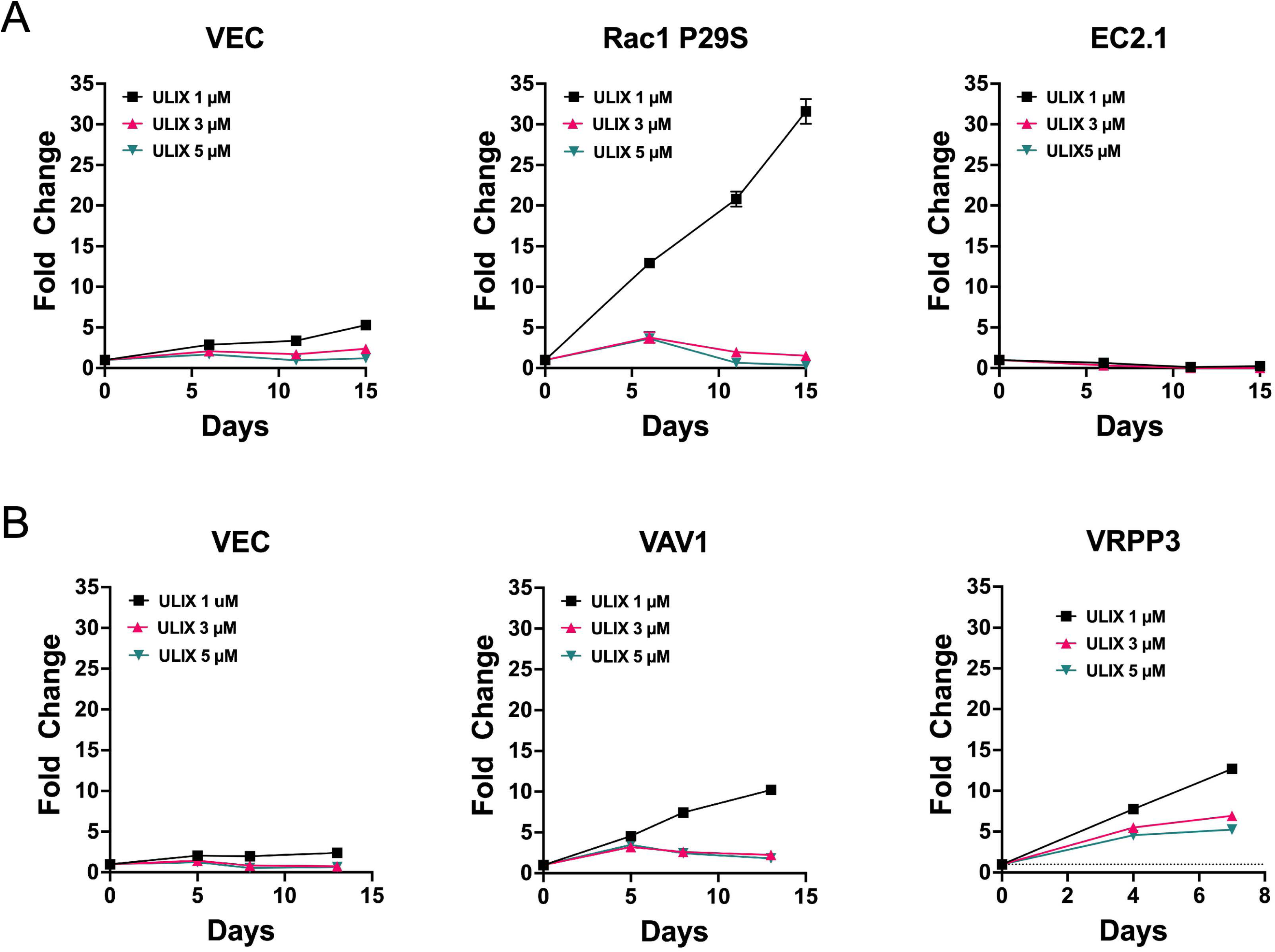
Rac-driven BRAFi-resistant melanoma cells are also partially resistant to ERK1/2 inhibition. **(A)** Vector control A375 cells (**VEC**), A375 cells expressing the Rac1 P29S mutant (**Rac1 P29S**), or A375 cells harboring a spontaneous BRAF truncation (**EC2.1**) were treated with 1 µM, 3 µM, or 5 µM ERK inhibitor, ulixertinib, and cell growth was measured over time via resazurin assay. Rac1 P29S cells were able to proliferate in 1 µM ulixertinib much better than vector control cells. However, EC2.1 cells were even more sensitive to ulixertinib than were vector control cells. **(B)** In a separate experiment, vector control A375 cells (**VEC**), VAV1 over-expressing cells (**VAV1**), or the Rac1 N92I mutant A375 subline (**VRPP3**) were treated with 1 µM, 3 µM, or 5 µM ulixertinib (**ULIX**) and cell growth was measured over time via resazurin assay. Both VAV1 overexpressing and VRPP3 cells were able to proliferate in 1 µM ulixertinib much better than vector control cells.

### Rac1-driven resistance mechanisms can be highly pleiotropic

We performed several additional experiments to investigate how Rac1 pathway signaling contributes to BRAFi/MEKi-treatment resistance, focusing on the A375 VRPP3 cells, which represent a spontaneous, Rac1-driven resistance mechanism to BRAFi-treatment. Because they displayed a YAP/TAZ-TEAD activation gene expression signature, we attempted to knock down YAP1 and TAZ via RNAi in A375 VRPP3 cells. Knocking down either YAP1 or TAZ partially reduced the BRAFi-resistance of VRPP3 cells (**Fig. S7A**). Notably, knocking down expression of the more highly abundant TAZ protein resulted in upregulation of the YAP1 protein (**Fig. S7B**). We have thus far been unable to obtain cells with strong, stable knockdown of both YAP1 and TAZ, suggesting that YAP1/TAZ may collectively be important for the Rac1-driven drug resistance mechanism in the A375 cell line.

In addition to signaling towards YAP1/TAZ, Rac1 is well known to activate alternate MAPK pathways involving Jun kinase and p38 MAPK. Consistent with this, signaling via Jun kinase and p38 MAPK appeared upregulated in A375 VRPP3 cells relative to parental cells or the A375 EC2.1 cells with a BRAF truncation, while signaling of ERK1/2 towards p90RSK appeared attenuated (**Fig. S8**), consistent with a reduced reliance of VRPP3 cells on ERK1/2, potentially via upregulation of other MAPK pathways. Therefore, we interrogated the A375 VRPP3 cell drug resistance mechanism with different combinations of BRAF, MEK, Jun kinase, and p38 MAPK inhibitors. Remarkably, only when the ERK1/2 MAPK, Jun kinase, and p38 MAPK pathways were simultaneously blocked were we able to completely inhibit the proliferation of the VRPP3 drug-resistant population (**Fig. S9**). While it remains to be determined how universal the pleiotropy of the Rac1-driven drug resistance mechanism is in BRAF V600-mutant melanoma, these data indicate that Rac1 signaling can drive MAPKi resistance via simultaneous parallel mechanisms that include the YAP/TAZ-TEAD, Jun kinase, and p38 MAPK pathways.

### Rac1-driven BRAFi/MEKi resistant melanoma cells respond to FAK inhibitors

Previously, we found that some Rac1-driven BRAFi-resistant melanoma cells showed a vulnerability to combination treatment with BRAFi and the selective Src kinase family inhibitor, saracatinib (Ref. (14) and **Fig. S4D**). However, saracatinib is no longer being developed clinically as a targeted therapy for cancer, and other Src inhibitors, such as dasatinib, are far less specific and inhibit many tyrosine kinases. Since Src kinases collaborate with focal adhesion kinase (FAK) to signal towards cell survival and proliferation, we hypothesized that FAK inhibitors (FAKi), which continue to be clinically investigated, might also show utility against Rac1-driven drug-resistant melanoma cells. Indeed, the combination of the BRAFi, vemurafenib, and the FAKi, defactinib, was effective not only in parental A375 (**Fig. S10A**), but also in the A375 VRPP sublines and A375 cells expressing VAV1 (**Fig. S10B-E**), as well as in 451Lu cells expressing the Rac1 P29S mutant (**Fig. S10F, G**). Since defactinib inhibits both FAK and the related kinase, Pyk2, we also used the FAK-selective inhibitor, PF 573228. This FAK-selective compound also blocked the growth of both A375 Rac1 P29S and VRPP3 cells (**Fig. S10H, I**).

An ongoing clinical trial is testing a novel RAF/MEK clamp, avutometinib (VS-6766) in combination with the FAK inhibitor defactinib for brain metastatic cutaneous melanoma (NCT06194929). We found that the combination of avutometinib + defactinib (or the combination of avutometinib and the FAKi VS-4718 used as a surrogate for defactinib in preclinical studies) showed activity against a variety of Rac1-driven BRAFi-resistant melanoma cell lines *in vitro* under both 2D (**Fig. 7A-J**) and 3D growth conditions (**Fig. 7K, L**). Strikingly, in the 3D growth conditions, cells expressing Rac1 P29S showed enhanced sensitivity to FAK inhibition alone compared to the vector control cells (**Fig. 7K, L**).

**Figure 7.**
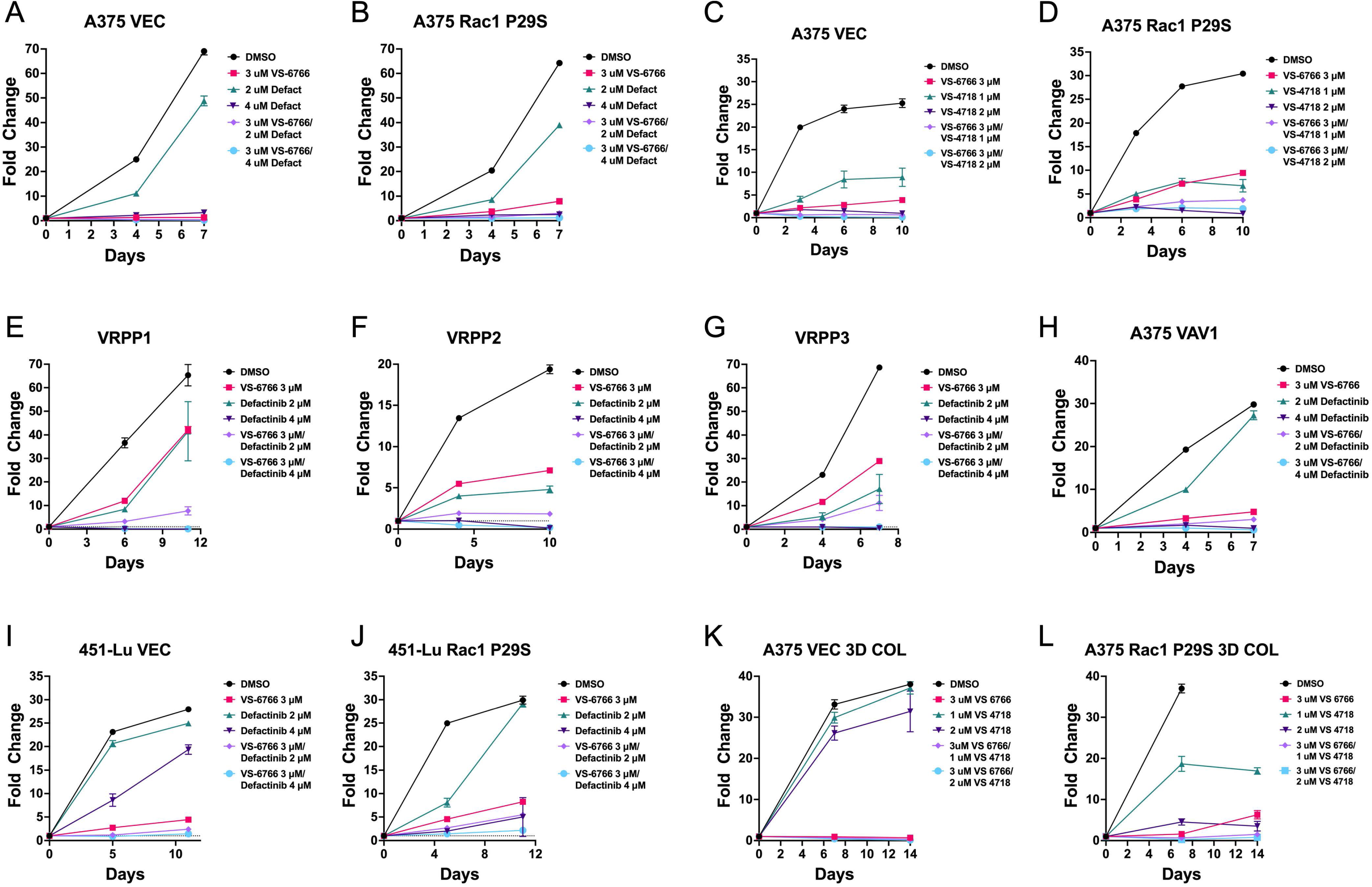
Rac1-driven BRAFi-resistant melanoma cells respond to combinatorial inhibition of RAF/MEK and FAK. (A-J) A375 and 451Lu cells, either vector control (VEC) or various Rac1-driven cell lines including Rac1 P29S-expressing cells, the A375 VRPP1-3 sublines, or A375 VAV1-expressing cells were treated with different combinations of the novel RAF/MEK clamp, VS-6766 (avutometinib) and the FAK inhibitors defactinib or VS-4718. Cell growth over time was measured via resazurin assay. While some cell lines displayed resistance to avutometinib alone, the combination of avutometinib and FAK inhibitors was able to control the growth of the cells. **(K&L)** Combined inhibition of RAF/MEK and FAK also controlled A375 Rac1 P29S cells growing on 3D collagen. Interestingly, cells expressing Rac1 P29S were also more sensitive to FAK inhibition as a monotherapy.

The above data highlighted the potential for avutometinib/defactinib as a treatment combination for BRAFi-resistant melanomas driven by Rac1, including melanomas harboring the Rac1 P29S mutation. To test this idea in an *in vivo* model, we used A375 Rac1 P29S cells, which are known to be resistant to BRAFi *in vivo* (13). Avutometinib plus VS-4718 showed substantial activity in controlling the growth of Rac1 P29S-mutant cells (**Fig. 8A**), significantly prolonging survival in this model (**Fig. 8B**).

**Figure 8.**
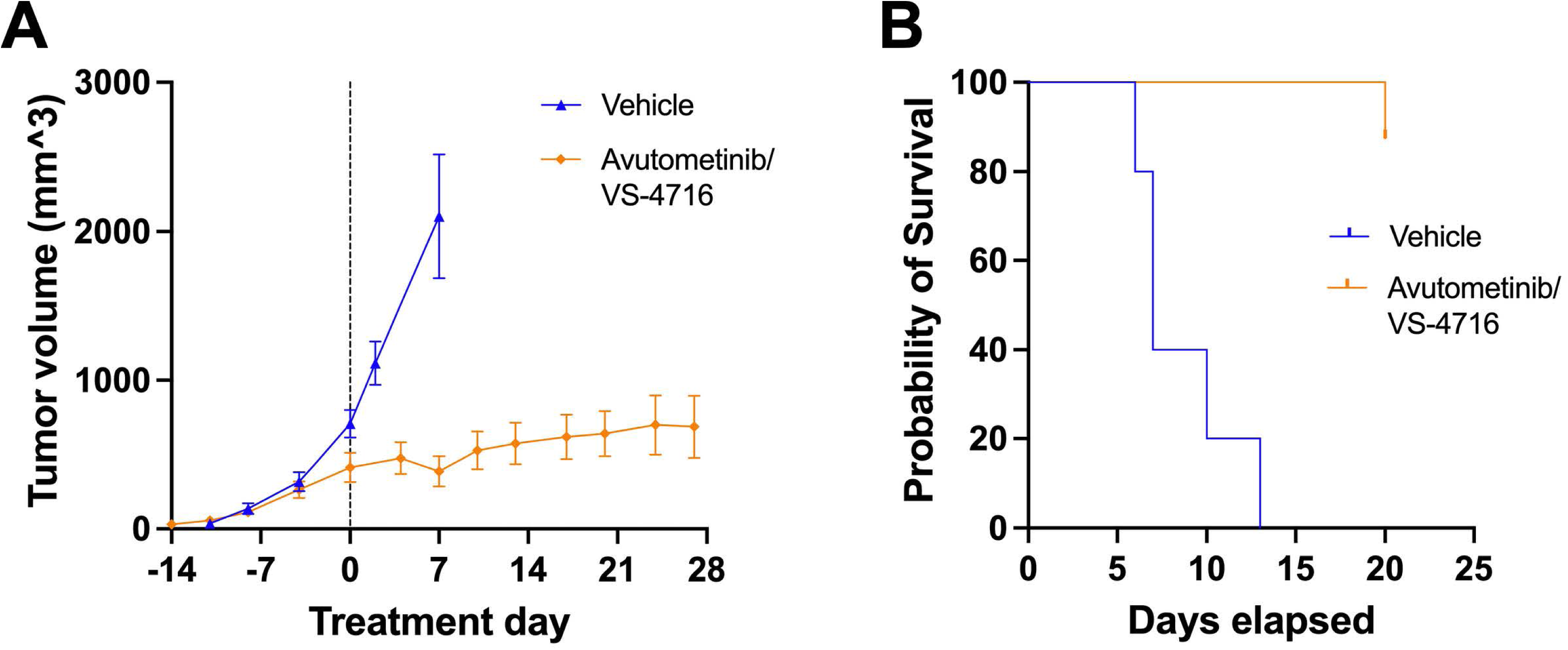
Inhibition of RAF/MEK and FAK show substantial activity in controlling the growth of Rac1 P29S-mutant cells. **(A)** 1 x 10^6^ A375 Rac1 P29S cells were implanted subcutaneously in the left and right flanks of 15 female nude mice (total of 30 tumors at the outset of the experiment). When tumors reached 500 mm^3^, mice were randomized and treated with vehicle control (5 mice) or avutometinib plus the FAK inhibitor VS-4718 (10 mice; treatment day 0 as indicated with the dashed line). **(B)** Kaplan-Meier analysis of survival of mice treated with vehicle control or avutometinib plus VS-4718. The drug combination significantly prolonged survival (P<0.0001).

## Discussion

BRAFi resistance mechanisms involving BRAF rearrangements, BRAF upregulation, growth factor receptor upregulation, and NF1 loss all converge on dimerization of mutant BRAF V600E, which normally signals as a drug-sensitive monomer (17, 18, 27, 28). Currently available FDA-approved BRAFi are unable to fully inhibit the activity of BRAF dimers, due to a dimerization-dependent conformational change that dramatically reduces the affinity of BRAFi for the second protomer in a BRAF dimer (18). In contrast to BRAF dimerization mechanisms of BRAFi resistance, the spontaneous resistance mechanisms driven by upregulation of Rac1 signaling are less well-understood. The uncertainty in precisely how Rac1 promotes BRAFi/MEKi resistance may reflect the fact that Rac1-driven resistance mechanisms can be pleiotropic, as we demonstrate here, involving reduced dependence on the ERK pathway and contributions from Jun kinase, p38MAPK, and YAP/TAZ-TEAD. The SRF/MRTF transcriptional complex can function as an additional effector of Rac1-driven BRAFi/MEKi resistance in some cell types (6), although SRF/MRTF inhibitor CCG-1423 did not resensitize our Rac1 P29S cells to BRAF inhibitors (data not shown).

Our results contrast somewhat with those of Mohan et al., who concluded that Rac1 P29S promotes ERK-independent, BRAFi-resistance that does not depend on Jun kinase, p38MAPK, or YAP/TAZ (9). This apparent contrast might be explained in part by the pleiotropy of Rac1 P29S-driven resistance mechanisms, in which several pathways make partial contributions to a collectively highly significant BRAFi-resistant phenotype. Such partial contributions may be less obvious in shorter term assays of cell proliferation, such as those used by Mohan et al. In addition, Mohan et al. highlighted yet another Rac1-driven resistance mechanism involving sequestration and inactivation of NF2 by Rac1-dependent lamellipodium formation.

PAK kinases, which are major effectors of Rac1, have been reported to promote BRAFi resistance at least in part by reactivating MAPK signaling via phosphorylation of MEK and CRAF(5, 7). Consistent with these prior reports, we found that inhibiting PAK signaling can partially abrogate Rac1-driven resistance. Using a PAK1-specific inhibitor, G-5555, we found that blocking PAK signaling can partially resensitize Rac1-driven cells to BRAF inhibition, without cytotoxic effects in the absence of BRAFi treatment. However, PAK phosphorylation of MEK kinase appears not to be mandatory for Rac1-driven resistance since our Rac1-driven cells were strongly resilient to MEK knockdown and inhibition, and a phosphomimetic mutation at the PAK-controlled S298 phosphorylation site on MEK did not promote BRAFi resistance. In addition, while PAK phosphorylation of CRAF may contribute to BRAFi resistance in some cells (5), we previously showed that VAV1-driven BRAFi resistance cannot be blocked by the pan-RAF inhibitor, LY3009120 (14). Regardless of which effectors function downstream of PAK to promote BRAFi/MEKi resistance, the use of PAK inhibitors in the clinic is likely to be limited by their on-target cardiotoxicity, which has even been observed for the PAK1-specific inhibitor G-5555 in preclinical studies (29).

The pleiotropic mechanism of Rac1-driven BRAFi/MEKi resistance, together with the difficulty of targeting the Rac1 effector PAK family kinases, creates a significant challenge for treating BRAF-mutant melanomas that develop Rac1-driven resistance to BRAF/MEK-targeted second line therapies after progression on front line immunotherapy. We have observed that selective inhibition of Src family tyrosine kinases can re-sensitize Rac-driven BRAFi-resistant cells to BRAF inhibitors (14, 22, 30) and **Fig. S4D**; however, selective Src kinase inhibitors are not currently being pursued clinically as anti-cancer therapies. Since Src kinases function together with FAK, we hypothesized that co-targeting BRAF/MEK and FAK with avutometinib (RAF/MEK clamp) plus defactinib (FAKi) would show activity against Rac1-driven drug-resistant melanoma cells. The *in vitro* and *in vivo* preclinical studies we present here provide strong support for this hypothesis.

Previous studies have placed Rac1 activation downstream of integrin-dependent FAK signaling (31, 32). Therefore, it is somewhat surprising that FAK inhibition can be highly effective in melanoma cells expressing constitutively activated Rac1, such as A375 Rac1 P29S and VRPP3 cells. Remarkably, on 3D collagen, A375 Rac1 P29S cells are even more sensitive to FAK inhibition than A375 vector control cells (**Fig. 7K, L**). In preliminary experiments, we found that FAK inhibition does not block PAK-controlled MEK S298 phosphorylation in A375 Rac1 P29S cells (data not shown), consistent with FAK-independent constitutive activity of the Rac1 P29S mutant. Nevertheless, avutometinib plus defactinib is effective at controlling the growth of Rac1 P29S cells, both *in vitro* and *in vivo*. Therefore, we currently favor the hypothesis that expression of constitutively active Rac1 can promote a FAK-dependent cellular phenotype in certain melanoma cells. The basis of such a FAK dependency remains to be determined.

Our results raise the priority to further define the cellular contexts in which FAK inhibition can overcome BRAFi/MEKi resistance. We previously observed that Rac1-Src signaling appears to drive BRAFi/MEKi resistance selectively in less differentiated melanoma cell lines such as A375 and 451Lu, which display downregulated MITF target gene expression and upregulated TEAD target gene expression (22). As described by Tsoi et al., melanoma cells responding to BRAFi treatment may pass through a phenotypic trajectory from a more differentiated state to a transitory state, to neural crest-like states and ultimately to an “undifferentiated” state (26), a phenotype that characterizes our A375 cell-derived VRPP BRAFi-resistant cells. The phenotypic starting point on this trajectory of cells undergoing BRAFi treatment may inform the extent to which Rac1 signaling can promote BRAFi adaptation and ultimately a BRAFi-resistant phenotype. An intriguing future direction will be to determine whether the differentiation status of Rac-driven cells is related to the extent to which they depend on FAK signaling.

Compared to MITF^low^ melanoma cells, such as A375 and 451Lu, cells beginning with a more differentiated, MITF^hi^ phenotype prior to drug treatment may be less able to utilize Rac1 signaling during drug adaptation (22). Interestingly, some melanoma cell lines with a more differentiated initial phenotype, such as UACC62 (22) and M229 (26), might tend to utilize RhoA signaling rather than Rac1 signaling as part of their adaptive response to BRAFi (33). Whether RhoA-driven BRAFi resistance is also responsive to FAK inhibition is another important open question.

A role for FAK in BRAFi resistance has previously been proposed, although not specifically in the context of Rac1-driven BRAFi resistance. FAK/Src signaling has been implicated in the initial adaptive response of BRAF-mutant melanoma cells to BRAFi treatments both *in vitro* (34) and *in vivo*, in response to an activated stroma (35). FAK has also been implicated in promoting a drug tolerant persister phenotype in a subset of human melanoma patient-derived xenographs (36). A key finding in our study is that FAK remains a relevant target in fully BRAFi-resistant melanoma cells driven by Rac1 activation. FAK signaling also promotes BRAFi resistance upon loss of AMBRA1 expression (37), but whether Rac1 is directly involved in this resistance mechanism is unknown. Based on a role for FAK in promoting melanoma brain metastasis (38), avutometinib plus defactinib is being tested in a clinical trial for brain metastatic cutaneous melanoma (NCT06194929). Our new results support the potential utility of this same combination against Rac1-driven, BRAFi-resistant disease.

## Materials and Methods

### Cell lines and Inhibitors

All cell culture reagents were obtained from Thermo Fisher Scientific (Waltham, MA), unless otherwise specified. A375 and 451Lu cells were obtained from the American Type Culture Collection (ATCC; Manassas, VA). GP2-293 cells (Takara Bio, San Jose, CA) and 293FT cells (Thermo Fisher Scientific, Waltham, MA) were used for production of retroviral and lentiviral particles respectively. All cell lines were cultured in DMEM, supplemented with 10% fetal bovine serum, penicillin/streptomycin, and L-glutamate. The derivation of BRAFi-resistant A375 subclones and populations has been previously described (14, 22). Briefly, A375 cells were exposed to 3 µM vemurafenib for 3-4 weeks until resistant cells appeared, either as rapidly growing colonies (EC1.1, EC2.1, EC7.1, ES2.2.1, ES2.2.2) or more slowly expanding drug-resistant populations (VRPP1-3). Rapidly growing colonies were isolated as clones using cloning cylinders, and VRPP1-3 were expanded and maintained as uncloned populations. The VRPP3 population acquired an activating Rac1 N92I mutation that went to fixation in the population (22). Pharmacological inhibitors used for cell-based assays in this study are listed in **Table S2**. Avutometinib and VS-4718 for *in vivo* studies were supplied by Verastem Oncology (Needham, MA).

### Transgene expression and RNA interference

Vectors used for gene expression and RNA interference (RNAi) are listed in **Table S2** in Supplementary Information. For gene transfer via the piggyBAC transposon/transposase system (39), cells were transfected with a 9:1 ratio (by µg) of transgene expression vector to piggyBAC transposase vector, then selected with 0.5 mg/ml G418 or 1 µg/ml puromycin beginning 48 hours later. For RNAi using the pSIREN vector backbone (Takara Bio, San Jose, CA), GP2-293 cells were transfected with a 5:1 ratio of shRNA vector to VSVG envelope protein expression vector. For RNAi using the pZIP-mCMV-ZsGreen-Puro vector backbone (Transomic Technlogies, Huntsville, AL), 293FT cells were transfected with a 4:3:1 ratio (by µg) of shRNA vector, to PAX retroviral structural vector, to VSVG envelope protein expression vector. At 24 and 48 hours after transfection, packaging cell media containing viral particles was harvested, filtered through a 0.45 µM syringe filter, and supplemented with 4 µg/ml polybrene before being used to transduce target cells. At 48 hours after the second round of transduction, target cells were selected with 0.3 mg/ml hygromycin or 1 µg/ml puromycin.

### Cell proliferation assays

To measure cell proliferation using resazurin (Thermo Fisher Scientific, Waltham, MA), 3,000-5,000 cells were plated per well in 4 wells per treatment group in 48 well plates. The next day (designated Day 0 of the growth assay), the relative cell number per well was measured by replacing the growth medium in each well with fresh growth medium containing 25 µg/ml resazurin sodium salt and incubated for 2 h at 37°C prior to reading fluorescence intensity at 590 nm using a BioTek Synergy HT plate reader (Agilent Technologies, Santa Clara, CA). Wells were then rinsed and refed with growth media containing various drug combinations (**Table S2**) or DMSO vehicle control. The relative number of cells per well was subsequently measured via resazurin at 3-4 day intervals and expressed as fold change relative to day 0.

To measure cell proliferation as population doubling versus time, 300,000 cells were plated per cell type/treatment condition in each of 3 wells in 6 well plates. Every 4-7 days, the cells were passaged, counted and the population doubling level (PDL) was calculated using the formula: PDLn = 3.32 (log Xt–log X0) +PDLn-1 (with Xt = cell number at that point, X0 = cell number used as inoculum and PDLn-1 = population doubling level at the previous passage). Vemurafenib treatment was renewed at each passage.

### Immunoblotting

Cells were rinsed twice with HBSM and lysed by scraping into Laemmli buffer. Protein concentrations of lysates were normalized using the Pierce 660 nm Protein Assay Reagent (Thermo Fisher Scientific, Waltham, MA) with SDS Neutralizer (G-Biosciences, Overland, MO) prior to SDS-PAGE and transfer to Immobilon-FL membranes (Millipore Sigma, Burlington, MA). Blots were blocked with Intercept TBS blocking buffer (LI-COR Biotechnology, Lincoln, NE) and then incubated with primary and secondary antibodies (**Table S2**). Blots were analyzed using a Odyssey blot imager (LI-COR Biotechnology, Lincoln, NE).

### RNAseq analysis of BRAFi-resistant cells

Parental A375 cells were cultured in DMEM, supplemented with 10% fetal bovine serum, penicillin/streptomycin, and L-glutamate. For 24- and 96-hour BRAFi-treated samples, vemurafenib was added to a final concentration of 3µM for the indicated intervals. All BRAFi-resistant A375 derivative cell lines were continuously cultured in 3µM vemurafenib. To extract total RNA, cells were trypsinized from three independent 10cm plates per condition, rinsed in PBS, and processed using the Direct-zol RNA Miniprep Plus kit (Zymo Research, Irvine, CA).

Sequencing libraries were prepared using the TruSeq Stranded mRNA kit (Illumina, San Diego, CA) and sequenced on a HiSeq-4000 device in 2 x 150bp paired-end format. Transcript pseudoalignment and quantification were performed with kallisto (40) using default settings, with Ensembl v94 transcript annotation as the reference transcriptome. Expression levels for each gene were obtained by summing transcript per million (tpm) values across all associated transcripts. These data are included as **Table S3**. All sequencing data are available in FASTQ format under GEO accession # GSE285131. Gene expression heatmaps and dendrograms were generated using Morpheus software from the Broad Institute (https://software.broadinstitute.org/morpheus). Gene level expression values were log-transformed using the log_2_(x+1) method, and unsupervised hierarchical clustering was performed based on the Euclidian distance metric.

For gene set enrichment analysis (GSEA), genes with maximum tpm values < 1 were excluded. VRPP1-3 samples were grouped and compared to all other samples, except for 24- and 96-hour vemurafenib-treated parental A375. Enrichment analyses were performed using GSEA (41, 42) desktop application version 4.0.1, with significance assessed based on 10,000 permutations of phenotype. Gene sets analyzed were “undifferentiated”, “neural crest-like”, “transitory”, and “melanocytic” from Table S3 of Tsoi et al. (26), as well as “YAP/TAZ Up” and “YAP/TAZ Down” from Table S3 of Kanai et al. (23).

### *In vivo* growth and treatment response of Rac1 P29S mutant cells

Female athymic nude mice (Strain # 007850; The Jackson Laboratory, Bar Harbor, ME) were injected subcutaneously in each flank with 1 x 10^6^ A375 cells expressing a firefly luciferase reporter and a RAC1 P29S transgene. Once average tumor size surpassed 500 mm^3^, one cohort of mice was treated with avutometinib (0.3 mg/kg once daily) plus VS-4718 (50 mg/kg twice daily) via oral gavage, while the other was gavaged with vehicle only on the same schedule. Vehicle for avutometinib was 5% DMSO, 10% hydroxypropyl-β-cyclodextrin in water. Vehicle for VS-4718 was 0.5% carboxymethyl cellulose, 0.1% Tween-80 in water. Mice were euthanized when tumor volume exceeded 2000 mm^3^. All animal studies were conducted using procedures approved and monitored by the Institutional Animal Care and Use Committee at the University of Iowa.

## Data Availability

The data generated in this study are available in the article and its supplementary files.

## Acknowledgements

Funding for this study was provided by the **Tad Agnew Foundation for Melanoma Research**. RNA sequencing experiments were performed at the Genomics Core Facility of the Holden Comprehensive Cancer Center in the University of Iowa Carver College of Medicine. The Holden Comprehensive Cancer Center is supported by the Center Core Grant P30 CA086862 awarded by the National Institutes of Health/National Cancer Institute.

## Author Contributions

JDR performed experiments, generated DNA constructs, help design and supervise in vivo experiments, performed RNAseq and bioinformatics analysis, and helped to draft and revise the manuscript. TAN developed cell lines, performed the MEK functional experiments, prepared figures and helped to draft the manuscript. AV developed cell lines, performed the Rac1 KD and Vav1 functional experiments, prepared figures, and helped to draft the manuscript. AAH, FAB, and NJF performed FAK inhibitor, Rac1 P29S, A375 VRPP3 experiments, prepared figures, and helped to draft the manuscript. RMP and ECH performed *in vivo* experiments and helped to prepare figures and draft the manuscript. MCJ helped develop and characterize spontaneously BRAFi-resistant A375 sublines, performed cell-based assays, and prepared figures for drafting the manuscript. SC and JAP contributed to the design of the *in vivo* experiments, provided key reagents and scientific input, and helped revise the manuscript. AJD and CSS supervised the study, designed experiments, performed experiments, analyzed data, and drafted and revised the manuscript.

## Ethics Declaration

All animal studies were conducted using procedures approved and monitored by the Institutional Animal Care and Use Committee at the University of Iowa.

### Competing Interests

SC and JAP are employees of Verastem Oncology

