## Supplementary figures and images for "A critical role of FAK signaling in Rac1-driven melanoma cell resistance to MAPK pathway inhibition"

### Supplemental Figures

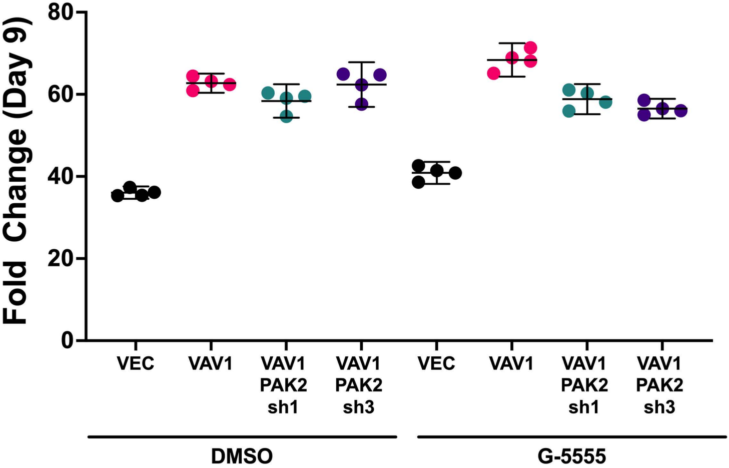

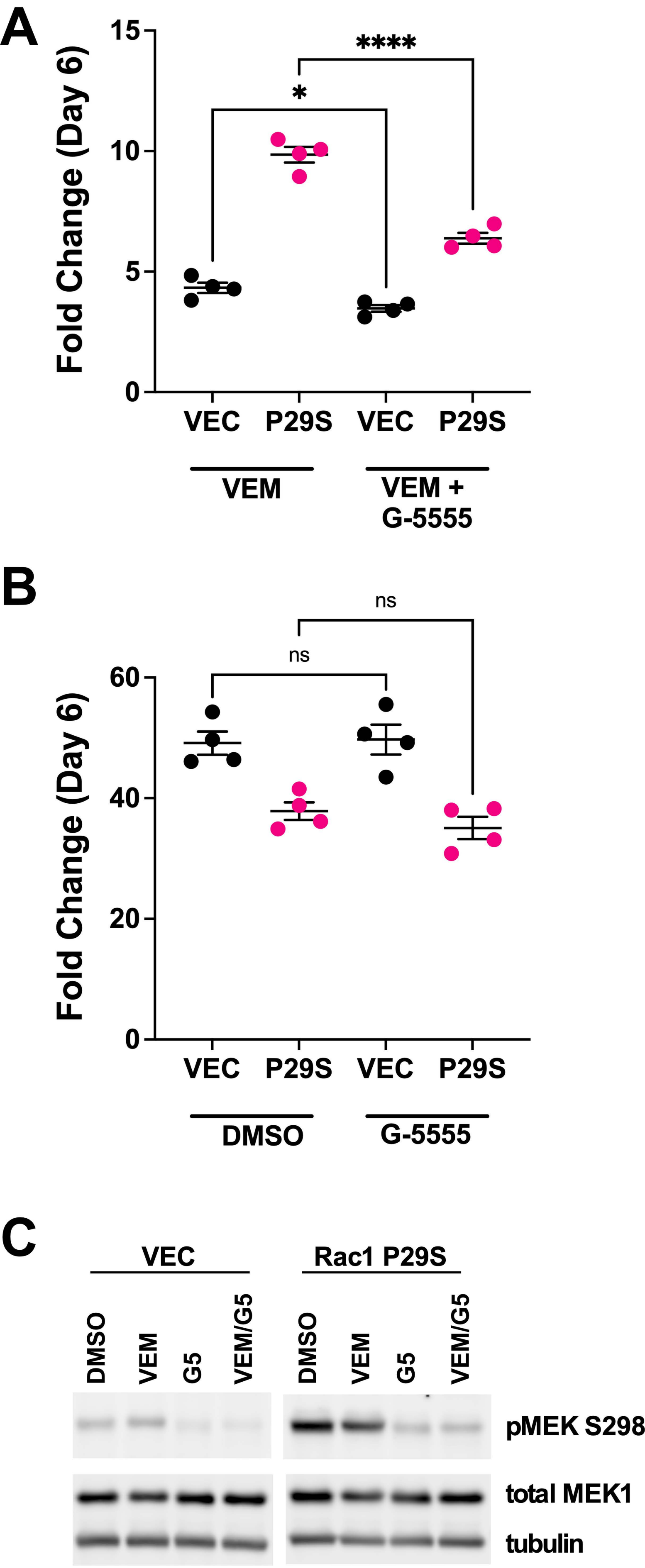

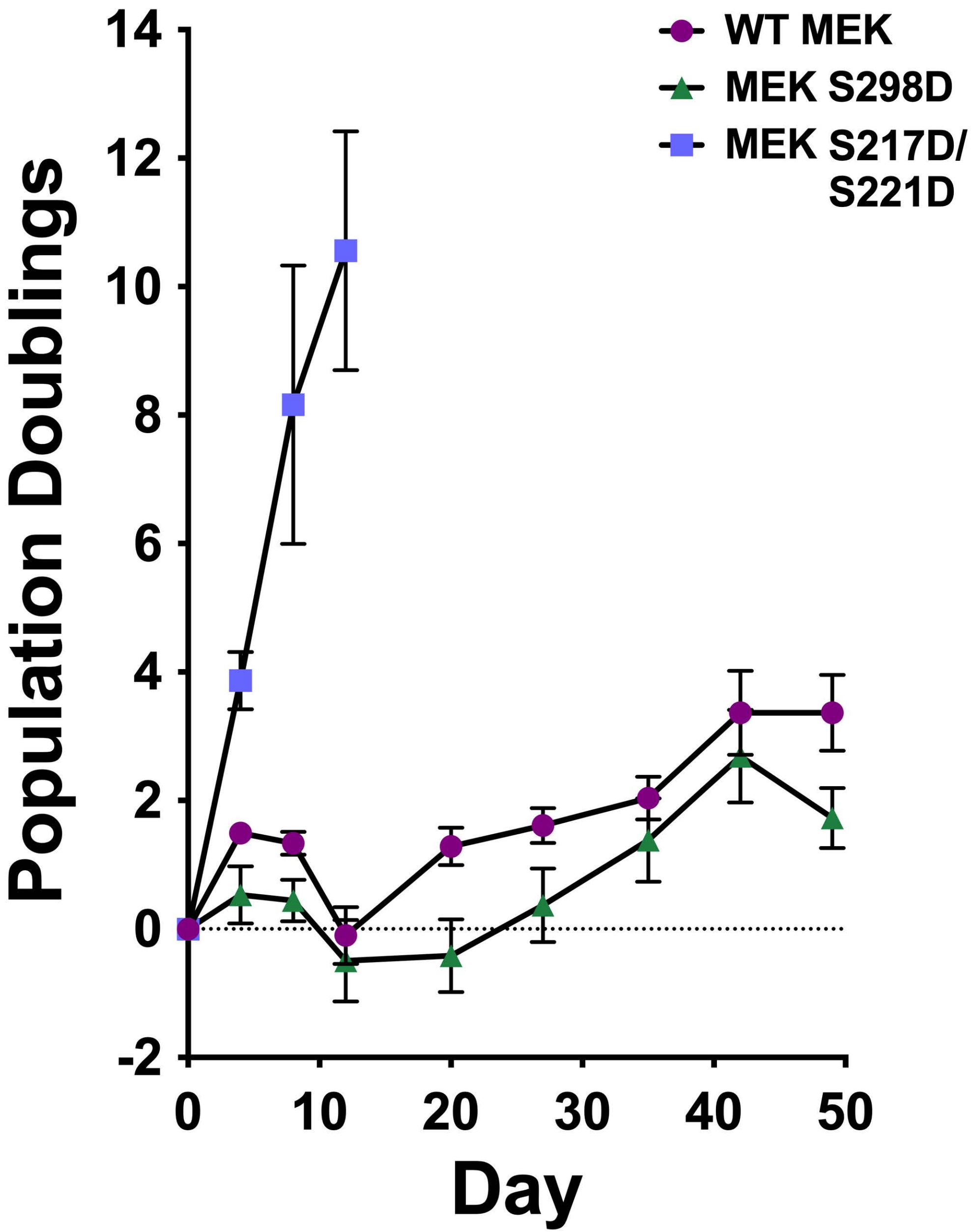

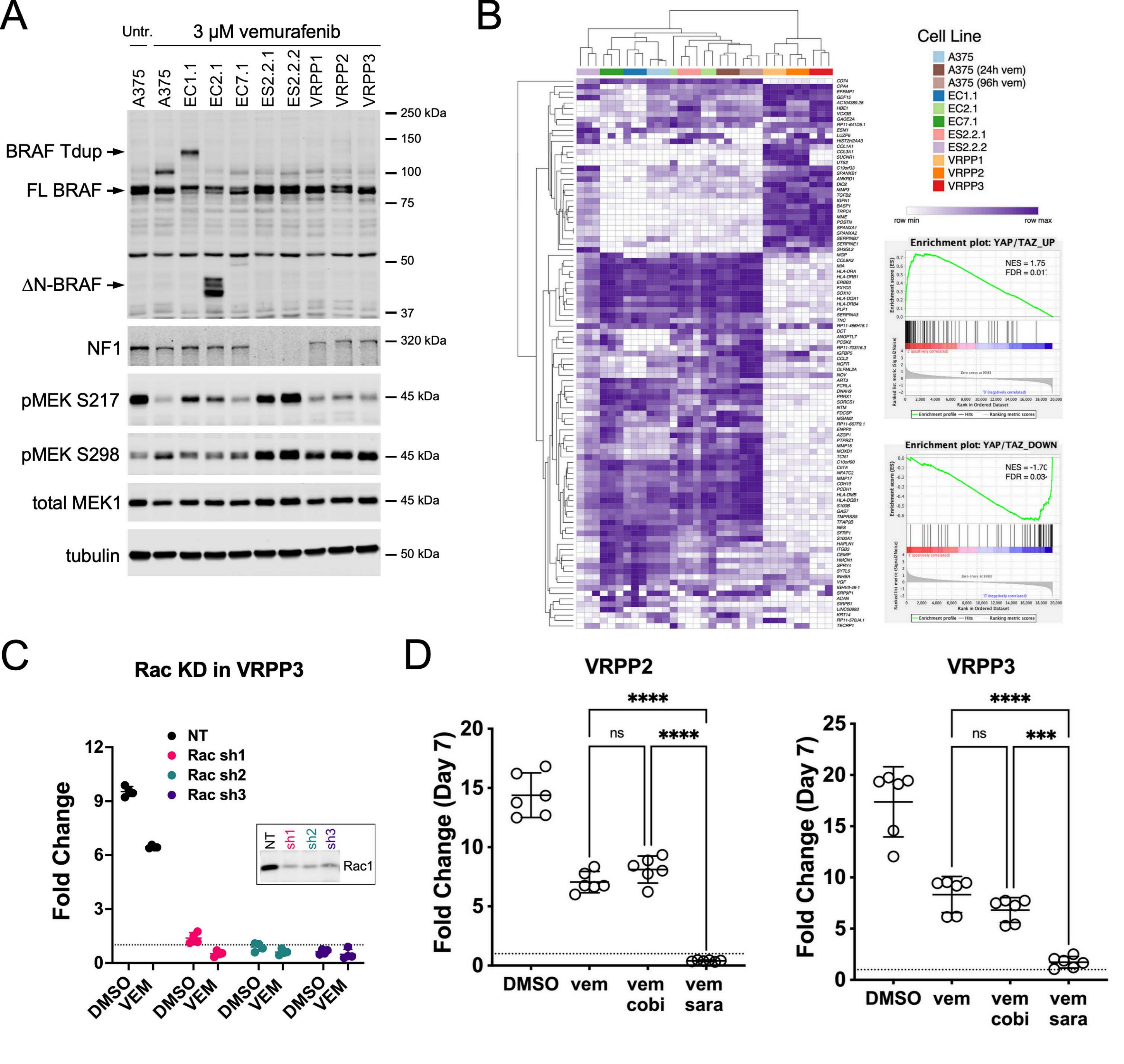

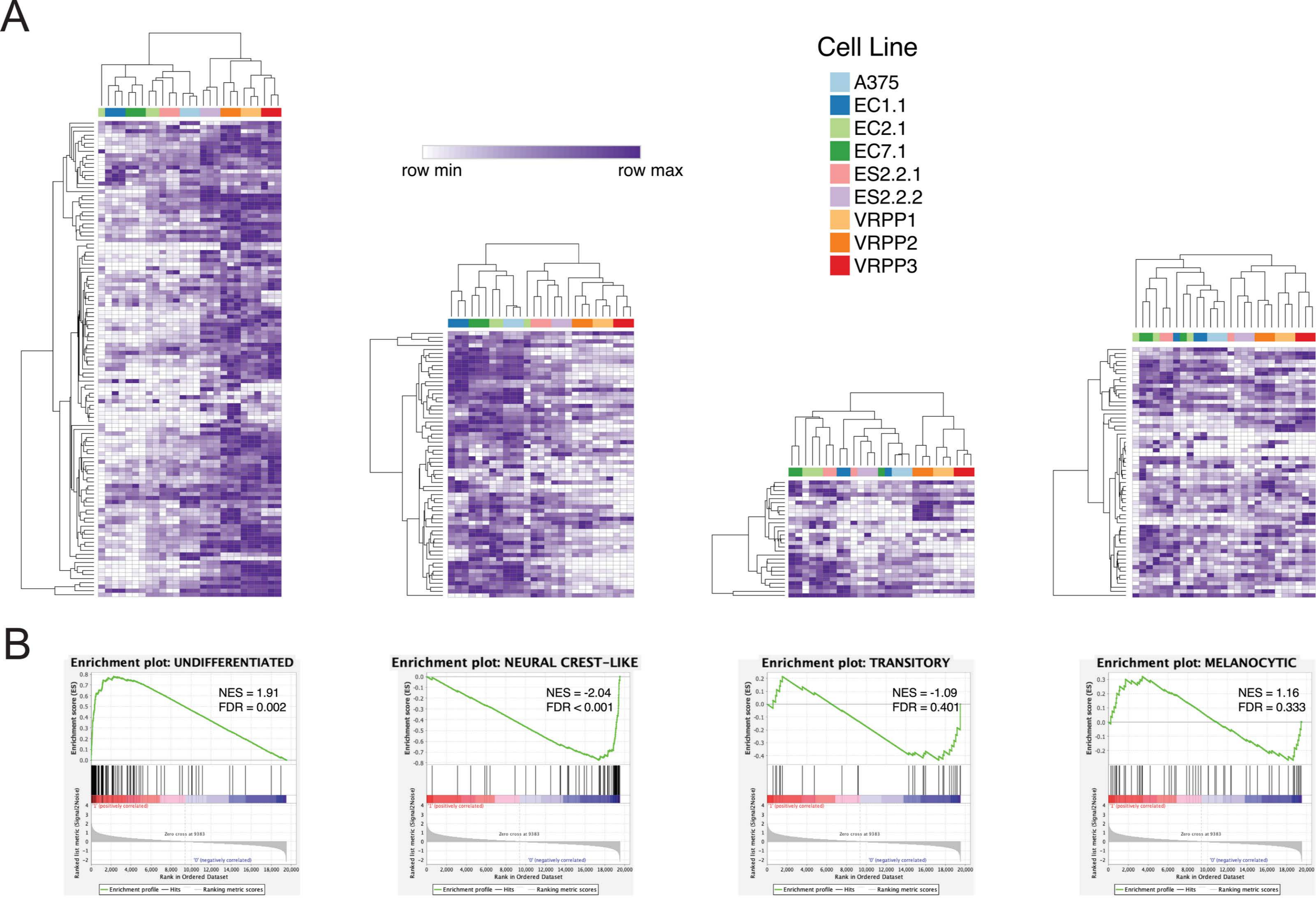

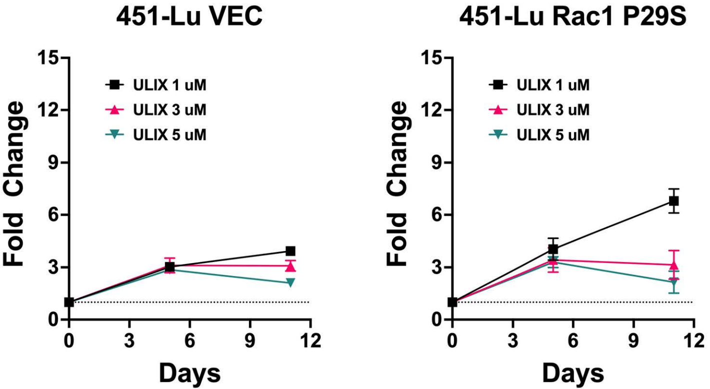

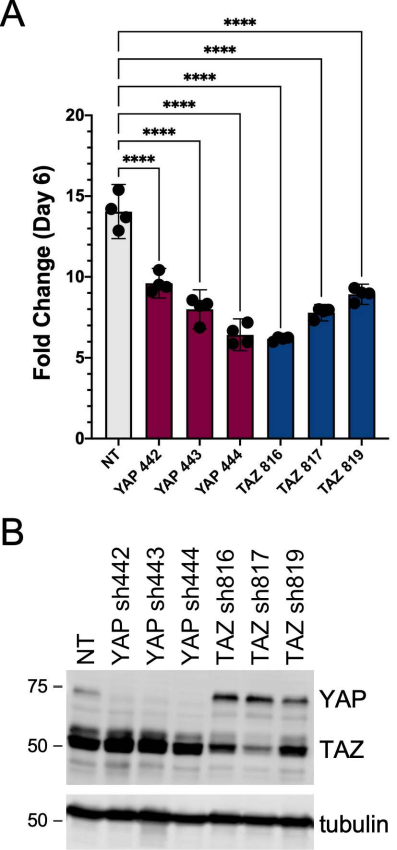

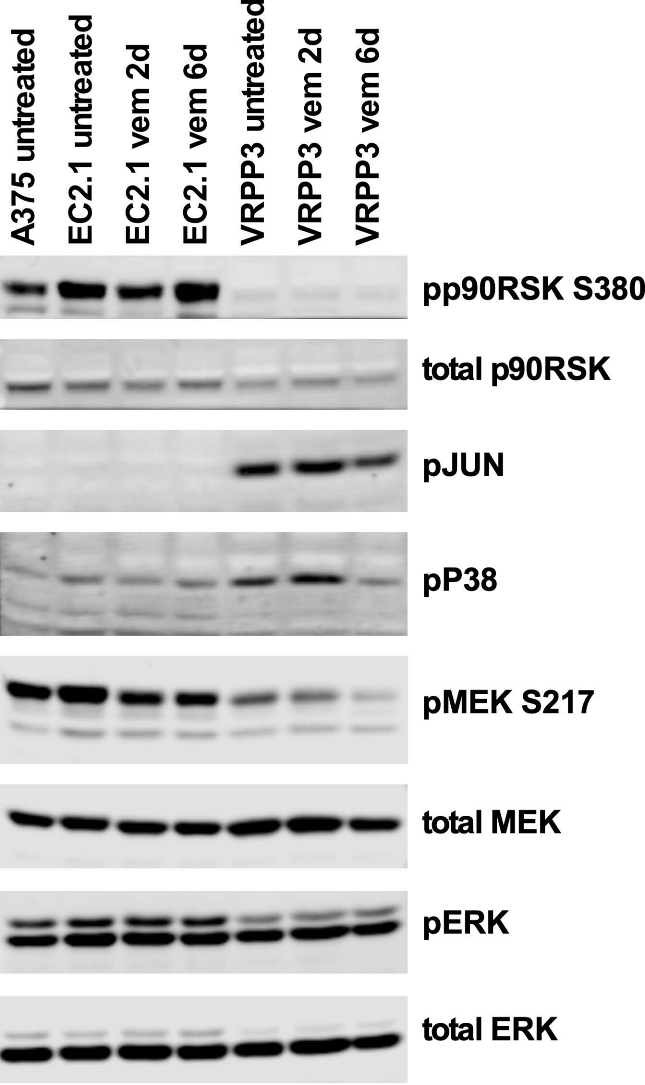

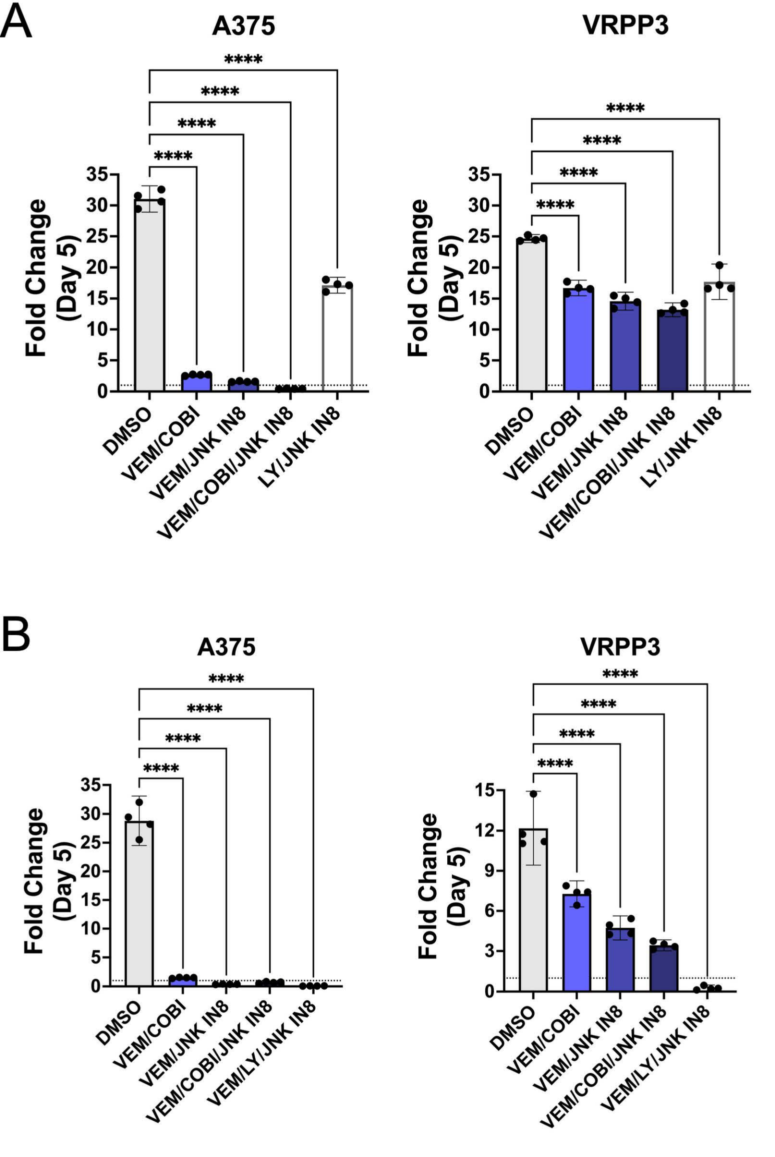

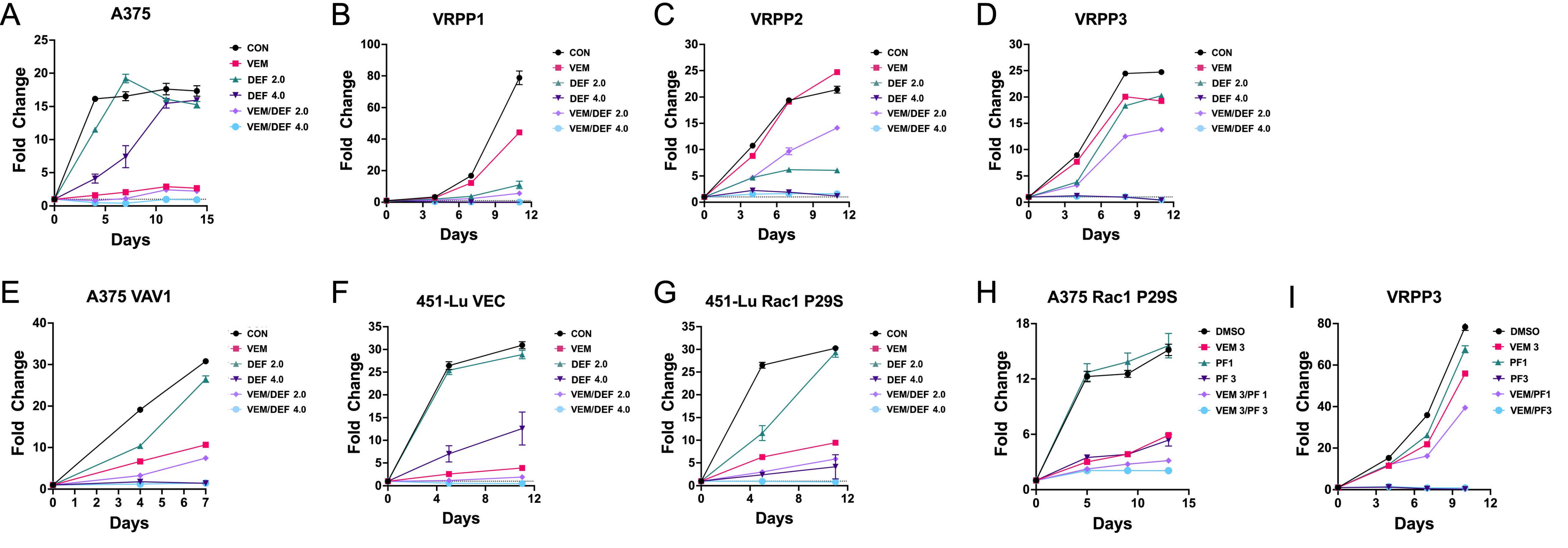
