## Supplemental Figure Legends for "A critical role of FAK signaling in Rac1-driven melanoma cell resistance to MAPK pathway inhibition"

**Supplementary Data**

**Figure S1. The PAK1-specific inhibitor G-5555 does not block the growth of Vav1-overexpressing cells on its own.** Vector control (VEC), Vav1 overexpressing cells (**VAV1**), or Vav1 overexpressing cells with either of two PAK2 shRNAs (VAV1 PAK2 sh1 and VAV1 PAK2 sh3) were treated with DMSO vehicle or 1 μM G-5555. Fold change in tumor cell number was measured on day 9 via resazurin assay.

**Figure S2. Rac1 P29S-driven BRAFi resistance in 451-Lu cells is partially PAK1-dependent. (A)** 451-Lu vector control cells (VEC), or cells harboring Rac1 P29S (P29S) were treated with 5 μM vemurafenib (VEM) or 5 μM vemurafenib plus 1 μM G-5555 (VEM + G-5555). Fold change in tumor cell number was measured on day 6 via resazurin assay. Treatment with vemurafenib plus G-5555 reduced cell proliferation compared to treatment with vemurafenib alone; *P<0.05, ****P<0.0001, ANOVA with Sidak's multiple comparison test. **(B)** The same cell lines as in (A) were treated with DMSO vehicle control or 1 μM G-5555 alone. **(C)** Lysates of vector control and Rac1 P29S cells were analyzed by SDS-PAGE and immunoblotting for phospho-MEK S298, total MEK1, and tubulin loading control. Rac1 P29S increased phosphorylation at PAK-controlled MEK S298, and this was largely abolished in cells treated with G-5555.

**Figure S3. A MEK BRAF phosphomimetic, but not a MEK PAK kinase phosphomimetic is sufficient to drive BRAFi resistance**. A population doubling versus time experiment showed that enforced expression of a MEK1 S217D/S221D mutant, a phosphomimetic at the BRAF-controlled serine-217 and 221 phosphorylation sites, promoted rapid proliferation in the presence of 3 μM vemurafenib. In contrast, enforced expression of a MEK1 S298D mutant, intended to mimic phosphorylation of MEK1 at the PAK kinase-controlled serine-298 phosphorylation site, failed to promote proliferation in the presence of 3 μM vemurafenib any better than enforced expression of wild type MEK1.

**Figure S4. Characteristics of spontaneously BRAFi-resistant A375 sublines. (A)** Lysates of parental A375 cells (A375) and A375 cells that acquired spontaneous BRAFi resistance by a variety of different mechanisms were analyzed by SDS-PAGE and immunoblotting. Spontaneous resistance mechanisms identified included tandem duplication of the BRAF kinase domain (BRAF Tdup) in EC1.1, BRAF N-terminal truncation (ΔN-BRAF) in EC2.1, loss of

RasGAP NF1 in ES2.2.1 and ES2.2.2 and increased signaling towards the PAK-controlled pMEK S298 site in the absence of the other mechanisms (VRPP1-3). Cells were either untreated, or treated with 3 μM vemurafenib, as indicated. **(B)** Gene expression was analyzed via RNA seq for parental A375 cells treated or not with 3 μM vemurafenib compared to each of the BRAFi resistant populations maintained under vemurafenib treatment. Unsupervised hierarchical clustering was performed based on levels of the top 100 variably expressed genes across all samples. The analysis revealed that the VRPP sublines display a distinct gene expression profile, characterized in part by increased expression of TEAD1/AP1 target genes. Statistical significance of this YAP/TAZ pathway enrichment was assessed by gene set enrichment analysis (GSEA) **(C)** As described in the text, the VRPP3 subline was found to have acquired a Rac1 N92I mutation. Knockdown of Rac1 in VRPP3 via three different retrovirally delivered shRNAs strongly inhibited growth under both control (DMSO) conditions and in the presence of 3 μM vemurafenib (VEM). Inset shows strong knockdown of Rac1 in each of the sublines. **(D)** VRPP2 or VRPP3 cells were treated with DMSO vehicle, 3 μM vemurafenib (vem), 3 μM vemurafenib plus 3 nM cobimetinib (vem/cobi), or 3 μM vemurafenib plus 2 μM saracatinib (vem/sara). Fold change in cell number was measured on day 7 via resazurin assay. Both cell lines were resistant to vemurafenib and cobimetinib, but were strongly blocked by the combination of vemurafenib and saracatinib; ***$P<0.001$, ****$P<0.0001$, ANOVA with Sidak's multiple comparison test.

**Figure S5. VRPP subline gene expression indicates transition from neural crest-like to undifferentiated state. (A)** Unsupervised hierarchical clustering was performed on RNA seq data from parental A375 cells and BRAFi resistant sublines based on gene sets identified by Tsoi et al. to represent four distinct stages of melanoma cell differentiation. Across these gene sets, VRPP sublines 1-3 clustered together. **(B)** Gene set enrichment analysis (GSEA) revealed strong, statistically significant enrichment for higher expression of “undifferentiated” genes and lower expression of “neural crest-like” genes in VRPP lines, as compared to all other samples.

**Figure S6. Rac1 P29S promotes resistance to ERK inhibition in 451-Lu cells.** 451-Lu vector control (VEC) and Rac1 P29S cells were treated with 1, 3, or 5 μM ulixertinib, and fold change in cell growth was measured over time via resazurin assay. Rac1 P29S expressing cells were partially resistant to 1 μM ulixertinib treatment.

**Figure S7. BRAFi resistance of A375 VRPP3 cells is partially dependent on YAP and TAZ expression. (A)** VRPP3 cells expressing a non-targeting shRNA (NT), or three different shRNAs each targeting YAP or TAZ were treated with 3 μM vemurafenib. Fold change in cell number was measured on day 6 via resazurin assay. All 6 shRNAs targeting YAP or TAZ partially suppressed the growth of the cells in vemurafenib; ****P<0.0001, ANOVA with Sidak's multiple comparison test. (B) Analysis of lysates of NT control cells and YAP or TAZ knockdown cells vis SDS-PAGE and immunoblotting showed effective knockdown of YAP and TAZ in cells with the YAP or TAZ shRNAs. In the TAZ knockdown cells, YAP was significantly upregulated compared to the level observed in the NT control cells. A tubulin loading control confirmed equal loading in each lane.

**Figure S8. Analysis of different MAP kinase signaling pathways in A375 EC2.1 and VRPP3 cells.** Lysates of A375 parental cells, EC2.1 cells with a BRAF truncation, and VRPP3 cells with a Rac1 N92I mutation were analyzed by SDS-PAGE and immunoblotting. Cells were untreated or treated with 3 μM vemurafenib for 2 or 6 days. Phospho-p90 RSK S380, an ERK1/2 target, was maintained in the EC2.1 cells, but not strongly expressed in VRPP3 cells regardless of drug treatment. In contrast, phospho-JUN and phospho-P38 MAP kinase appeared increased in VRPP3 cells. Phospho-MEK S217, a BRAF-controlled site, was strongly maintained in EC2.1 cells, but reduced VRPP3 cells. Both phospho- and total ERK1 appeared somewhat reduced in VRPP3 compared to the other sublines. Overall, the differences in MAP kinase signaling in VRPP3 appeared consistent with a reduced reliance on ERK signaling in those cells.

**Figure S9. The A375 VRPP3 subline utilizes multiple MAP kinase pathways to resist drug treatments. (A)** Parental A375 or the VRPP3 subline were treated with DMSO vehicle, or different combinations of BRAF inhibitor vemurafenib (3 μM; VEM), MEK inhibitor cobimetinib (3 nM: COBI), Jun kinase inhibitor JNK-IN-8 (3 μM) or p38 MAP kinase inhibitor LY2228820 (5 μM; LY). **(B)** A separate experiment from that shown in (A), using an additional treatment in which BRAF, p38 MAP kinase, and Jun kinase were all inhibited simultaneously. Only this triple combination was able to completely suppress the drug resistant phenotype of the VRPP3 cells.

**Figure S10. Inhibition of FAK can block the BRAFi resistance of Rac1-driven melanoma cells.** (**A-G)** A375 parental cells (A), VRPP1-3 sublines (B-D), A375 Vav1 overexpressing cells (E), or 451-Lu vector or Rac1 P29S cells (F-G) were treated with DMSO vehicle (CON), 3 μM vemurafenib (A375 cells) or 5 μM vemurafenib (451-Lu cells) (VEM), 2 μM defactinib (DEF 2.0),

4 μM defactinib (DEF 4.0), or vemurafenib combined with defactinib (VEM/DEF 2.0 and VEM DEF 4.0). Fold change in cell growth was measured over time via resazurin assay. Co-treatment with defactinib blocked resistance to vemurafenib and in some cases cells responded to defactinib treatment alone. **(H&I)** In a similar experiment, defactinib was replaced with the FAK-specific inhibitor PF 573228 at 1 μM or 3 μM to treat A375 Rac1 P29S or VRPP3 cells. PF 573228 was able to cancel the BRAFi resistance of both cell lines, supporting FAK as the critical target of defactinib, which can also inhibit PYK2.
