## Supplemental Table S2 for "A critical role of FAK signaling in Rac1-driven melanoma cell resistance to MAPK pathway inhibition"

**Table S2. Pharmacological reagents, vectors, and antibodies used in this study**

| **Pharmacological reagents** | | | | |
| --- | --- | --- | --- | --- |
| **Compound** | **Target** | | | **Source** |
| Vemurafenib | BRAF V600E | | | MedChemExpress |
| G-5555 | PAK1 | | | Targetmol |
| Cobimetinib | MEK1/2 | | | MedChemExpress |
| Ulixertinib | ERK1/2 | | | Targetmol |
| Avutometinib (VS-6766) | MEK1/2 | | | Targetmol (*in vitro* studies)  Verastem (*in vivo* studies) |
| Defactinib | FAK/Pyk2 | | | Targetmol |
| VS-4718 | FAK/Pyk2 | | | Targetmol (*in vitro* studies)  Verastem (*in vivo* studies) |
| PF 573228 | FAK | | | MedChemExpress |
| Saracatinib | SRC family kinases | | | MedChemExpress |
| JNK-IN-8 | JNK | | | MedChemExpress |
| LY2228820 | P38 MAPK | | | MedChemExpress |
| **Vectors** | | | | |
| **Backbone** | **Insert** | | | |
| PB-EF1α-MCS-PGK-G418 | VAV1 | | | |
| PB-EF1α-MCS-PGK-G418 | empty | | | |
| PB-EF1α-MCS-PGK-Hygro | VAV1 | | | |
| PB-EF1α-MCS-PGK-Hygro | empty | | | |
| PB-EF1α-MCS-PGK-Puro | Rac1 P29S | | | |
| PB-EF1α-MCS-PGK-Puro | MEK1 wild type-myc | | | |
| PB-EF1α-MCS-PGK-Puro | MEK1 S217D, S221D-myc | | | |
| PB-EF1α-MCS-PGK-Puro | MEK1 S298D-myc | | | |
| PB-EF1α-MCS-PGK-Puro | empty | | | |
| pZIP-mCMV-ZsGreen-Puro | Rac1 sh1 (5’-CAAGGAGATTGGTGCTGTAAAA-3’) | | | |
| pZIP-mCMV-ZsGreen-Puro | Rac1 sh2 (5’-CCAAGAAGATTATGACAGATTA-3’) | | | |
| pZIP-mCMV-ZsGreen-Puro | Rac1 sh3 (5’-CGAATATATCCCTACTGTCTTA-3’) | | | |
| pZIP-mCMV-ZsGreen-Puro | YAP1 sh442 (5’-AGAAAGCTTTCTTACATGGTT-3’) | | | |
| pZIP-mCMV-ZsGreen-Puro | YAP1 sh443 (5’-CACATCGATCAGACAACAACA-3’) | | | |
| pZIP-mCMV-ZsGreen-Puro | YAP1 sh444 (5’-AGGTGATACTATCAACCAAAT-3’) | | | |
| pZIP-mCMV-ZsGreen-Puro | TAZ sh816 (5’-TCCGGAGGACTTCCTCAGCAA-3’) | | | |
| pZIP-mCMV-ZsGreen-Puro | TAZ sh817 (5’-CACATAGAAAAAATCACCACA-3’) | | | |
| pZIP-mCMV-ZsGreen-Puro | TAZ sh819 (5’-CCGGAGGACTTCCTCAGCAAT-3’) | | | |
| pZIP-mCMV-ZsGreen-Puro | Non-targeting | | | |
| pSIREN-RetroQ-Puro | shMEK1 (5’-CCGCAGAGAGAGCAGATTTGA-3’) | | | |
| pSIREN-RetroQ-Hygro | shMEK2 (5’-CTCAAAGACGATGACTTCGAA-3’) | | | |
| pSIREN-RetroQ-Hygro | PAK2 sh1 (5’-AGAAATTTCTCCTCCATCTGA-3’) | | | |
| pSIREN-RetroQ-Hygro | PAK2 sh3 (5’-CAGGAGGTTGCTATCAAACAA-3’) | | | |
| pSIREN-RetroQ-Puro | Non-targeting | | | |
| pSIREN-RetroQ-Hygro | Non-targeting | | | |
| **Antibodies** | | | | |
| **Antibody** | **Clone** | **Catalog No.** | **Source** | |
| Rabbit anti-BRAF V600E | RM8 | MA5-24661 | Thermo Fisher Scientific | |
| Mouse anti-ERK1/2 | L3F12 | 4696 | Cell Signaling Technology | |
| Rabbit anti-p-ERK1/2 T202/Y204 | D13.14.4E | 4370 | Cell Signaling Technology | |
| Mouse anti-MEK1 | 61B12 | 2352 | Cell Signaling Technology | |
| Rabbit anti-MEK2 | polyclonal | 9125 | Cell Signaling Technology | |
| Rabbit anti-p-MEK S298 | D1P9E | 98195 | Cell Signaling Technology | |
| Rabbit anti-p-MEK S217/221 | 41G9 | 9154 | Cell Signaling Technology | |
| Rabbit anti-NF1 | D7R7D | 14623 | Cell Signaling Technology | |
| Rabbit anti-p90RSK | 32D7 | 9355 | Cell Signaling Technology | |
| Rabbit anti-p-p90RSK S380 | D3H11 | 11989 | Cell Signaling Technology | |
| Rabbit anti-PAK2 | C17A10 | 2615 | Cell Signaling Technology | |
| Rabbit anti-p-c-JUN S73 | D47G9 | 3270 | Cell Signaling Technology | |
| Rabbit anti-p-P38 T180/Y182 | D3F9 | 4511 | Cell Signaling Technology | |
| Mouse anti-Rac1 | 102 | 610651 | BD Biosciences | |
| Mouse anti-tubulin | 12G10 |  | DSHB | |
| Rabbit anti-Vav1 | polyclonal | HPA001864 | Sigma | |
| Rabbit anti-YAP/TAZ | D24E4 | 8418 | Cell Signaling Technology | |
